# Distinct senescent β-cell senotypes differentially drive islet aging and dysfunction

**DOI:** 10.64898/2026.05.25.727705

**Authors:** Kanako Iwasaki, Hui Pan, Jonathan Dreyfuss, Maya Jackson, Sergii Domanski, Dylan Baker, Priscila Carapeto, Christopher Cahill, Sandra Le, Francesko Hela, Juliana Alcoforado Diniz, Giray Naim Erylmaz, Fan Wu, Pei-Hsun Wu, Bofei Yu, Denis Wirtz, Sara Espinoza, Alejandro Pena, Francisco G. Cigarroa, Gregory Abrahamian, Jillian L. Woodworth, Peter Adams, Duygu Ucar, Jeffrey H. Chuang, Vesna D Garovic, James L. Kirkland, Tamar Tchkonia, Nicolas Musi, George A. Kuchel, Paul Robson, Cristina Aguayo-Mazzucato

## Abstract

Biological aging greatly impacts the body’s ability to handle glucose, and represents a major risk factor the development and progression of type 2 diabetes (T2D). Nonetheless, despite advances in cellular senescence research and the development of new senolytic therapies, the heterogeneity of cellular senescence in the human endocrine pancreas, as well as its roles in normal aging, remains to be elucidated at the single-cell level.

Here, we performed single-cell–resolved spatial proteomics and transcriptomics on intact pancreas from 26 donors (ages 20–80) and multiplexed single-cell RNA sequencing and functional assays on dispersed islets from 14 donors (ages 34–69). We identify two discrete SnC subpopulations distinguished by relative expression of CDKN1A and CDKN2A.

*CDKN1A*⁺ senescent cells (SnCs) exhibit loss of β-cell identity, impaired insulin secretion, and a proinflammatory SASP associated with increased islet immune infiltration. In contrast, *CDKN2A*⁺ SnCs retain transcriptional identity and functional competence, with lower inflammatory signaling.

Together, these findings identify heterogeneous and functionally divergent senotypes in the human pancreas, distinguishing an adaptive (*CDKN2A*⁺) from a maladaptive (*CDKN1A*⁺) senescence program, thus providing a mechanism-guided framework for senescence-targeted therapies in T2D.

## INTRODUCTION

Senescent Cells (SnCs) and the senescence-associated secretory phenotype (SASP) are heterogeneous, with diverse roles in health and disease. Characterization and detection of distinctive SnCs subpopulations, or “senotypes”, is a priority in developing targeted and more effective therapies. To this end, the study, detection, and mapping of SnCs across the human lifespan aim to advance diagnostic and therapeutic approaches that improve human health [1].

The human pancreas is a complex organ with distinct endocrine hormone-secreting cells (α, β, δ, and γ) in the islets of Langerhans whose primary function is to regulate blood glucose levels. The biological relevance of SnCs in islets has been linked to Type 1, Type 2, and monogenic diabetes [2]. However, it is unknown whether these populations are homogeneous and whether they have specific adaptive or maladaptive roles.

SnCs in the endocrine pancreas have reported antagonistic roles with both positive [3] and negative effects on β-cell function [4], and evidence for both protective [5] and disease-promoting roles [6] in diabetes pathophysiology. Overexpression of p16 in both mouse and human islets, was shown to improve glucose-stimulated insulin secretion (GSIS) and drive expression of key β-cell genes such as *Insulin* and *Mafa* [3], suggesting the existence of adaptive SnCs. Meanwhile, a β-cell SnC signature of senescence-associated β-galactosidase (SA-βGal^+^) cells, demonstrated overall downregulation of identity genes (including *Insulin* and *Mafa*) and upregulation of disallowed genes which are normally suppressed in this cell population [4]. In humans, SA-βGal^+^ β-cells increased with age, body mass index and Type 2 Diabetes, suggesting a maladaptive phenotype. Results are further confounded by the use of different mouse models [7], and limited access to high-quality human tissue samples.

We hypothesized that previous contradictory findings in pancreatic β-cell senescence could be explained by the existence of distinct SnCs subpopulations within human islets. We further proposed that these subpopulations display unique molecular, spatial, and functional programs that differentially contribute to endocrine pancreatic aging and dysfunction. We define these distinct senescent states as senotypes.

Systematic analysis of SnCs led to the identification of two senotypes defined by *CDKN1A* (encodes p21) and/or *CDKN2A* (encodes p16) expression with distinct distributions. Cell-specific senescence marker and senescence-associated secretory phenotype (SASP) signatures were developed for all endocrine cell types. The secretome from *CDKN1A^+/^ CDKN2A-* and *CDKN1A^+/^ CDKN2A^+^* senescent β-cells was immune regulatory in nature and linked to higher islet immune infiltration when compared to the *CDKN1A^-/^ CDKN2A^+^* SASP. Transcriptional senotype profiling revealed loss of cellular identity in *CDKN1A^+/^ CDKN2A-* SnCs associated with a dysfunctional response to secretory stimuli, as shown by a lack of insulin secretion in response to high glucose concentrations. On the other hand, *CDKN1A^-/^CDKN2A^+^* SnCs retained their transcriptional identity and function.

Together, our findings reveal distinct adaptive and maladaptive β-cell senotypes in the human endocrine pancreas. We identified an adaptive CDKN1A⁻/CDKN2A⁺ state and a maladaptive, immune-regulatory CDKN1A⁺/CDKN2A⁻ senotype linked to impaired islet function and tissue inflammation. These results establish a framework for understanding how SnCs heterogeneity contributes to pancreatic aging and provide a foundation for senotype-specific therapeutic interventions.

## RESULTS

### β-cell senescence as a pathognomonic feature of human aging

The identification of senotypes in the endocrine pancreas leveraged spatial transcriptomics Xenium, and Visium with spatial proteomics CODEX, in whole pancreas from heart beating donors across the lifespan (**Fig. 1A**). This was complemented with proteomic analysis of the secretome and the evaluation of isolated pancreatic islets using both static and dynamic methods to assess glucose stimulated insulin secretion (GSIS) as a gold standard for β-cell function (**Supplemental Tables 1**).

**Figure 1.**
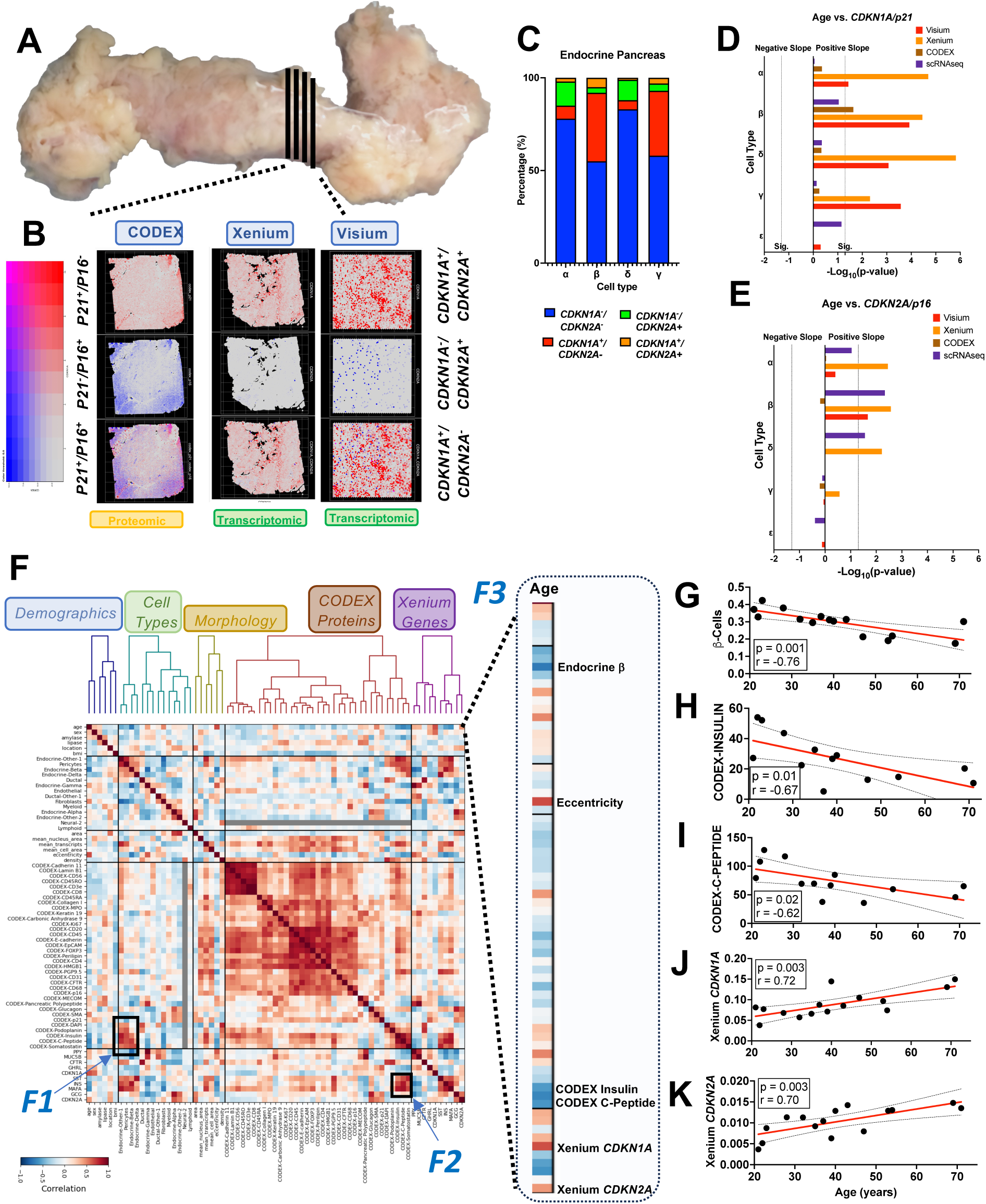
Senotype identification in human endocrine pancreas and age-correlated variables highlight β-cell senescence as a key feature of human aging. **(A)** Representative pancreas from a 67-year-old male donor with a BMI of 29 and no history of diabetes. Total weight 131.6 g. **(B)** Spatial feature plots and blended image plots to visualize co-expression of *CDKN1A* and *CDKN2A* in Visium and Xenium processed specific samples, and p21 AND p16 for selected CODEX samples. Color threshold code expresses the levels of *CDKN1A/*p21 (red) and *CDKN2A*/p16 (blue). **(C)** Percentage of 2 senotypes (*CDKN1A^+^/ CDKN2A*^-^, *CDKN1A^-^/ CDKN2A*^+^), double positive, and double negative in the human endocrine cell types from scRNASeq data obtained from 13 donors; Correlation between age and *CDKN1A* **(D)** and *CDKN2A* **(E)** in endocrine pancreas cell types from all platforms. The is the log(10) of the p value, and the dotted line represents a significant p-value of 0.05. Negative and positive slopes are represented in opposite sides of the y axis. **(F)** Heatmap of correlation of variables across 32 tissues. The variables are separated into categories: demographic, cell type morphology, CODEX protein and Xenium genes calculated as a robust median across islets in each tissue. Within each group, variables are clustered hierarchically. Dark red and dark blue squares show high (*F1, F2*) and low Pearson correlation respectively. Gray squares correspond to values with undefined correlation. The inset highlights correlation of age and all other variables (*F3*). Age-related signal quantification in individual donors of β-cell proportion **(G)**, CODEX protein levels of INSULIN **(H)** and C-PEPTIDE **(I)** and Xenium gene levels of *CDKN1A* **(J)** and *CDKN2A* **(K)**. Mean+/-SD are plotted for each donor and red line represents average value per age group.

Senotypes were defined by the expression of SnC markers and effectors *CDKN1A* (encodes p21) and *CDKN2A* (encodes p16) on the following criteria: (1) their presence was detected in all cell types across pancreas - omic platforms (scRNASeq, Visium, Xenium and CODEX) (**Suppl. Fig. 1A-C**) and (2) their spatially distinct distributions at the transcriptomic and proteomic levels suggested discrete identities (**Fig. 1B**).

Each endocrine cell type was divided into a non-SnC *CDKN1A^-^/CDKN2A^-^* state and one of two following senotypes: *CDKN1A^+/^ CDKN2A^-^* and *CDKN1A^-/^ CDKN2A^+^* and a double positive *CDKN1A^+/^ CDKN2A^+^.* Senotype prevalence in the endocrine pancreas was studied using scRNASeq and indicated β- and γ- cells as being those with the highest percentage of SnCs (45% and 42% respectively) and *CDKN1A^+/^ CDKN2A^-^* as the dominant senotype (37% in β-cells and 35% in γ-cells) (**Fig. 1C, Suppl. Fig. 2A**). These percentages were significantly higher in scRNASeq studies when compared to spatial approaches where the pancreatic tissue was intact such as Xenium and CODEX (**Suppl. Fig. 2B**). Tissue manipulation prior to sequencing likely accounts for the increase in SnCs. Orthogonal testing of identified senotypes across different technological platforms mitigate this phenomenon.

Evaluation of senotypes across the human lifespan revealed a significant and positive correlation between *CDKN1A* transcription and chronological age in all endocrine cell types in Visium and Xenium platforms (**Fig. 1D**), while *CDKN2A* transcription was positively associated with age in α, β and δ cells by Xenium and β−cells by Visium (**Fig. 1E**). No significant correlations were found between either senotype and body mass index (BMI) or glycosylated hemoglobin (HbA1c) (**Suppl. Fig. 2C-F**) consistent with all donors being metabolically healthy.

The variables from 32 tissue samples (**Suppl. Table 1**) were separated into demographic variables, cell type morphology, CODEX protein, and Xenium genes and analyzed for correlational strengths with Pearson coefficient (r) including all cell types found in and immediately around the islets except acinar (**Fig. 1F**). High correlation clusters revealed known associations between protein levels of INSULIN and C-PEPTIDE with both β-cells and transcription of hallmark genes such as *INSULIN* and *MAFA* (**Fig. 1F, inset *F1* and *F2***). Identification of features with the highest negative or positive correlations with age (**Fig. 1F inset *F3***) revealed the strongest negative associations with the proportion of β-cells in islets (**Fig. 1G**) which were paralleled by decreased protein content of the identity hormone INSULIN (**Fig. 1H**) and intragranular C-PEPTIDE (**Fig. 1I**). Both senotype identity genes revealed an age-dependent increase in *CDKN1A* (**Fig. 1J**) and *CDKN2A* (**Fig. 1K**) transcription.

These results point to the accumulation of SnCs in the endocrine pancreas across the human lifespan and highlight β-cell senescence, based on increased *CDKN1A* and *CDKN2A* expression and decreased β-cell identity and abundance, as a key feature of human aging.

### Anatomical determinants of endocrine senotype distribution

Whole pancreas were sectioned into pancreatic portions (inferior and superior head, body, and tail) following anatomical features highlighted by surgical guidelines [8]. Visium spatial transcriptomic analysis revealed the presence of all senotypes in every pancreatic region (**Fig. 2A**) and in every endocrine cell type (**Suppl. Fig. 3A-F**). The inferior head was characterized by a predominance of *CDKN1A^+/^ CDKN2A^-^* γ-cells that was not observed in other regions of the pancreas. This unique distribution and cell-type enrichment is consistent with the distinct embryological origin of the inferior head from the ventral endoderm bud, while the superior head, body, and tail originate from the dorsal endoderm bud. Senotype gradients were observed for *CDKN1A^+/^ CDKN2A^-^* β-cells (**Suppl. Fig. 3C**) with a higher proportion in the superior head that decreased towards the tail. This gradient was mirrored by *CDKN1A^-^ /CDKN2A^+^*α-cells (**Suppl. Fig. 3B**). The opposite gradient dynamic was observed for *CDKN1A^+/^ CDKN2A^-^* δ-cells (**Suppl. Fig. 3D**), with the highest percentage in the tail.

**Figure 2.**
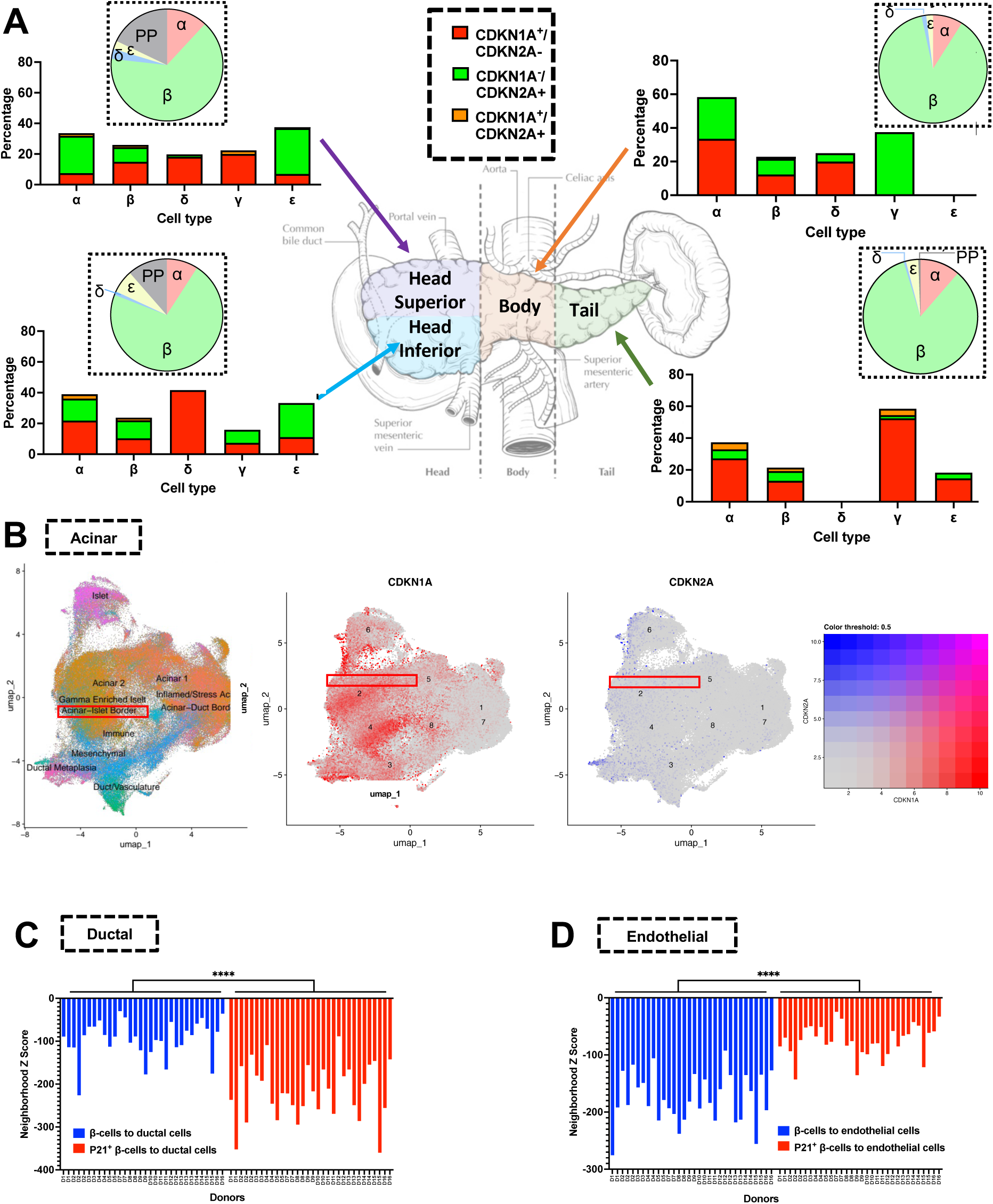
Spatial distribution of endocrine senotypes and effects on islet composition. (**A**) Senotype proportions of endocrine pancreas cell types per pancreatic proportion from Visium data following the surgical pancreatic guidelines [8] (n=10 donors, 40 samples). Pie charts represent average proportion of different endocrine cell types in islets from different pancreatic regions. (**J**) UMAP of Visium data colored by cell types and senotype visualization. Color threshold code expresses the levels of *CDKN1A* (red) and *CDKN2A* (blue) (**K**) Neighborhood proximity analysis between ducts and p21^-^ β-cells (blue bars) and p21^+^ β-cells (red bars). (**L**) Neighborhood proximity analysis between endothelial cells outside islets and p21^-^ β-cells (blue bars) and p21^+^ β-cells (red bars).

Analysis of the anatomical relationship of senotypes with other anatomical structures in the pancreas included spatial senotype mapping in the whole pancreas which revealed a *CDKN1A* enrichment in the Acinar-Islet Border (**Fig. 2B**). Spatial neighborhood proximity analysis was performed between p21^+^ β-cells and ductal and endothelial cells. The distance between ducts and p21^+^ β-cells was greater than to non-SnCs (**Fig. 2C**) supporting the known role of these structures in endocrine cell differentiation and development [9]. Endothelial cells outside the islet (**Fig. 2D**) were significantly closer to SnC p21^+^ β-cells suggesting a pro-senescence vascular-endocrine communication.

### Age-dependent remodeling of islet architecture and senotypes

Islet size, architecture, and composition are important for islet function with metabolic implications. Age-related changes in islet architecture were queried through calculation of an eccentricity score (1-Centrality) and revealed islet elongation with age (**Fig. 3A, B, B’**).

**Figure 3.**
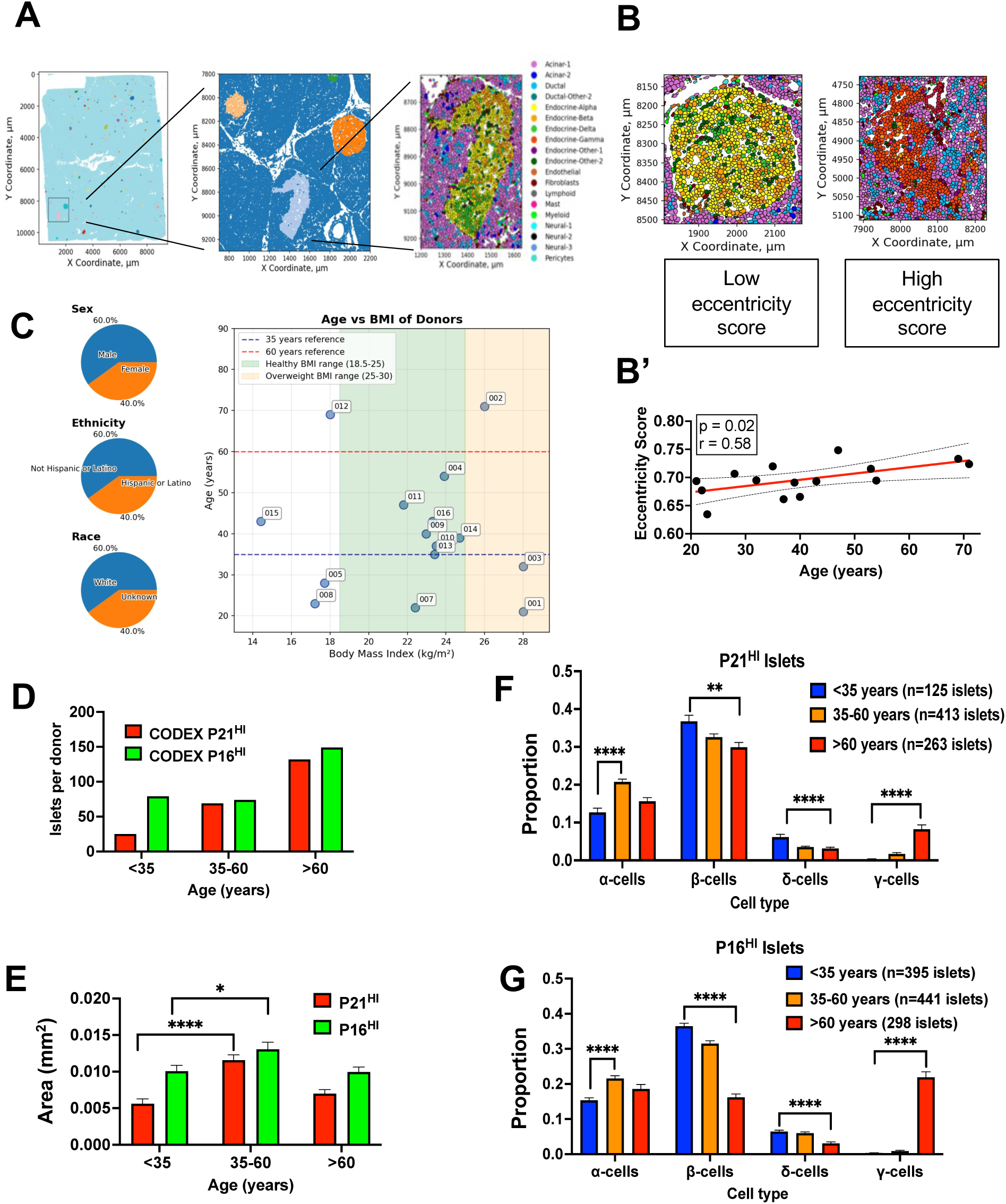
**(A)** Tissue section JDC-WP-008-b spatial layout highlights islets shaded with random colors on light blue background. A small, boxed tissue region is magnified in the middle inset featuring four islets, where the largest islet is elongated (high eccentricity) and is further magnified to show its cell composition including endocrine and non-endocrine cell types. The tissue surrounding the islet is primarily composed of acinar cells. (**B**) Age-related islet eccentricity (1-Centrality) quantification with representative islet inset for different scores (**B’**). Linear correlation between mean eccentricity score values and age. (**D**) Demographic characteristics and health profiles of 15 whole pancreas donors are presented. The plot illustrates the relationship between donor age and body mass index (BMI), with reference lines at 35 years (blue dashed) and 60 years (red dashed). Shaded regions indicate the healthy BMI range (18.5-25 kg/m², green) and overweight range (25-30 kg/m², tan). Individual donors are identified by their identification numbers. The donor cohort is approximately evenly distributed by sex (60% male, 40% female) and ethnicity (60% Non-Hispanic/Latino, 40% Hispanic/Latino). Race is reported as 60% White and 40% Unknown. Most donors fall within the healthy BMI range and the middle-aged group. (**D-G**) Hierarchical clustering of pancreatic islets reveals donor-specific endocrine composition patterns across different age groups and selection criteria. Islets from young (<35, n=5), middle-aged (35-60, n=6), and older donors (>60, n=2) were selected based on high expression of CODEX-p16 or CODEX-p21 (normalized fluorescence intensity >2.0) and hierarchically clustered using Pearson correlation with average linkage based on the proportions of α, β, δ, and γ cells. (**D**) Quantification of the number of p21^HI^ (high intensity: HI) and p16^HI^ islets per donor in each age category. (**E**) Average area of p21^HI^ and p16^HI^ islets in different age groups. Mean+/-SEM; *p<0.05; (**F**) Islet cell composition data of p21^HI^ and (**G**) p16^HI^ islets. Mean+/-SEM; *p<0.05. The number of islets analyzed for each age range was as follows: <35 years- 520 islets; 35-60 years- 854 islets; >60 years- 561 islets.

Association of senotypes with islet composition was analyzed in a cohort of 15 whole pancreases that represented lifespan, gender, and ethnicities over a wide BMI range (**Fig. 3C**). Islets from young, middle-aged, and older donors were selected based on high expression of CODEX-p16 or CODEX-p21 and interrogated for area and cell composition. The results demonstrated that islets cluster primarily by endocrine composition rather than by donor, with significant heterogeneity in senescence marker expression and cell type distribution across all age groups and selection criteria (**Suppl. Fig. 3G**). Quantification of number of islets per donor revealed that in donors younger than 35 years of age p16^+^ islets were the predominant senotype while the number of p21^+^ islets increased with age (**Fig. 3D**). Examination of changes in the area of SnC islets showed that islet size peaked in the 35-60 age range for both senotypes (**Fig. 3E**). Predominance of *CDKN1A^-/^ CDKN2A^+^* larger islets in younger individuals is consistent with an adaptive senotype.

Age altered the composition of SnC islets with high proportions of p21+ (**Fig. 3F**) or p16^+^ (**Fig. 3G**) cells. SnC p21^HI^ islets decreased the proportions of β and δ cells accompanied by an increase in γ-cells. A peak in the number of α-cells in SnC p21^HI^ islets was observed in donors 35- to 60-year-old group. Changes in SnC p16^HI^ islets followed a similar pattern with a more accentuated increase in γ-cells supporting an increase in cell complexity with age.

These results reveal a non-random distribution of SnC islets in the pancreas with significant age-related changes in size, structure, and cell composition marked by decreased β and δ-cells. The reduction of β-cells with age, coupled with changes in paracrine effects of somatostatin from δ-cell and glucagon from α-cells, can explain some of the reported age-related decline in β-cell function [10] and underscores the relevance of SnCs in the aging of the endocrine pancreas.

### Senotypes and senescence biomarkers

Given the heterogeneity of senescent cells (SnCs), no single biomarker has been universally validated across all senotypes. Accordingly, high-plex profiling approaches are recommended to accurately define SnC identity and diversity [2]. However, the limited scalability of these technologies underscores the need to identify robust, accessible biomarkers that can distinguish senotypes.

To understand whether specific SnC biomarkers preferentially correlated with specific senotypes, scRNASeq, CODEX platform and direct testing of dispersed human were queried for a panel of commonly used markers of SnC, at the mRNA, protein and enzymatic level.

Biomarkers for the hallmarks of senescence were assessed across senotypes at the scRNA-seq level (**Fig. 4A**). *CDKN1A*^+^ β-cells were enriched for *TP53* and *RB1* cell cycle arrest genes, upregulation of antiapoptotic pathways *BCL2L1,* and *EFNB1, GDF*15 as a SASP factor and *TM4SF1* as a cell surface marker. *CDKN2A*+ cells upregulated the *ABL1* antiapoptotic pathways and displayed levels of *CD36* and *NOTCH1* as cell surface markers. Interestingly, double positive *CDKN1A/CDKN2A* β-cells bore a higher resemblance to *CDKN1A*^+^ cells.

**Figure 4.**
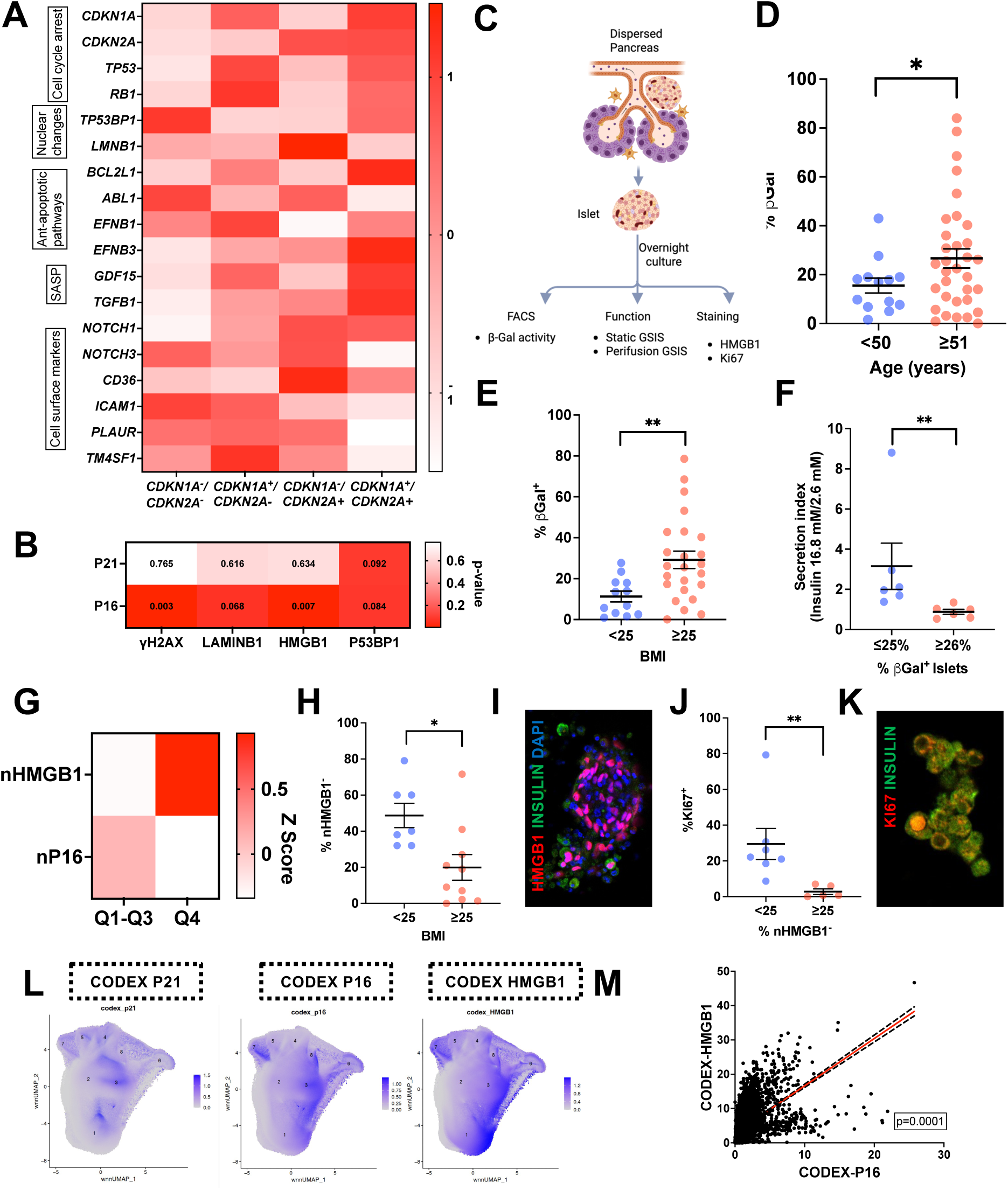
Senotypes and senescence biomarkers. (**A**) Heatmap of scRNASeq gene expression levels of hallmarks of senescence across b-cell senotypes. (**B**) Heatmap showing correlation between CODEX expression of p21 and p16 senotypes in β-cells with senescence biomarkers. Correlation p-values are included for each correlation. (**C**) Graphical abstract of workflow on islets obtained from human dispersed pancreas. Percentage of SA-βGal^+^ cells in dispersed whole pancreas (islet, acinar, and duct fractions plotted individually) from donor groups by age (**D**) and BMI (**E**). (**F**) Donor-matched %SA-βGal^+^ cells as assessed by flow cytometry with Secretion Index (Insulin secreted at 16.8 mM glucose/insulin secreted at 2.6 mM glucose during glucose stimulated insulin secretion (GSIS)). Mean+/-SEM, n=6 each, t-test, *p<0.05. (**G**) Heatmap of CODEX protein expression to emphasize a correlation between nuclear p16 (np16) and nuclear HMGB1 (nHMGB1). (**H, I**) Immunostaining quantification and representative image of HMGB1 staining in dispersed whole pancreas (islet, acinar, and duct fractions plotted individually) in 5 donors with different BMI; (**J**) Correlation of proliferating Ki67^+^ cells in dispersed pancreatic cells with senescent nHMGB1**^-^** cells; (**K**) Representative image of Ki67 staining. (**L**) Visualization of the expression of protein senescence markers p21, p16 and HMGB1 on UMAP plots from integrated CODEX and Xenium data. (**M**) Linear correlation of CODEX-HMGB1 and CODEX-p16 in islets from individual donors.

Additional biomarkers of SnCs were analyzed at the protein level using CODEX data in both p21 and p16 senotypes (**Fig. 3B**). DNA damage SnC biomarker p53BP1 positively associated with both senotypes. However, γH2AX, LAMINB1, and HMGB1 (high mobility group box 1) levels correlated with p16, but not p21, suggesting senotype specificity.

Direct testing of overnight cultured islets, ducts and acinar cells included evaluation of additional markers such as SA-βGal activitiy, staining of HMGB1 and KI67 to evaluate proliferation (**Fig. 3C**).

Senescence associated β-galactosidase (SA-βGal) activity, a well-known senescence enzymatic marker [11], was assessed in live pancreatic cells by flow cytometry and showed a significant increase with age (**Fig. 3D, Suppl. Fig. 4A**). This marker was also increased in pancreas from donors with a BMI >25 (**Fig. 3E**), which is consistent with previous reports of age and insulin resistance increasing β-cell senescence [4]. To understand whether this increase had functional consequences, the secretion index (fold change between insulin secreted at low and high glucose concentrations) was calculated and revealed a significant decrease in β-cell function in islets with a SA-βGal^+^ content of 26% or higher (**Fig. 3F**). These dynamics resemble those a maladaptive senotype which is dysfunctional and increases with age and states of insulin resistance.

HMGB1 is used as a biomarker of SnCs whose expression increases at the cellular level and undergoes subcellular redistribution by being extruded from the nuclei as SnC develops [12]. To further inquire the relationship between the HMGB1 as a SnC biomarker and p16^+^ islet cells, CODEX data were segmented to isolate the nucleus in β-cells. Nuclear levels of HMGB1 were divided into quartiles and interrogated for p16 levels. Lower levels of nuclear HMGB1 (nHMGB1, consistent with SnC), associated with higher levels of nuclear p16, supporting senotype specificity (**Fig. 3G**). nHMGB1^-^ SnCs were lower in pancreas from donors with a higher BMI (**Fig. 3H, I**) (in contrast to SA-βGal^+^) and did not correlate with age (**Suppl. Fig. 4B**), consistent with an adaptive senotype. As expected for a SnC biomarker, higher levels of nHMGB1^-^ associated with decreased proliferation levels (**Fig. 3J, K**).

Further senotype specificity was studied by visualizing the spatial distribution of p21, p16 and HMGB1 on whole pancreas UMAPS obtained from CODEX data (**Fig. 3L**). The expression pattern of HMGB1 followed that of p16 but not p21. Quantification of this correlation confirmed that β-cells with higher content of HMGB1 protein had also higher levels of p16 (**Fig. 3M**).

These results show that SnCs biomarkers at the protein level can be used to track specific senotypes in the endocrine pancreas and will be useful in the study of these subpopulations. HMGB1 expression correlated with the *CDKN2A^+^* an adaptive senotype and while SA-βGal^+^ islet cells track the dynamics of a maladaptive senotype.

### Senotype-specific Senescence Associated Secretory Phenotype (SASP)

SnCs secrete a bioactive SASP with important effects in neighboring cells. Given the existence of senotypes in the endocrine pancreas, we inquired whether they had specific SASP profiles. Our proteomic analysis of conditioned media from human islets ([13], GSE162521) was cross referenced with scRNASeq to define whether known secreted SASP factors were preferentially transcribed by a given senotype (**Suppl. Fig. 5**). The top 20 factors from *CDKN1A^+^/CDKN2A^-^*, *CDKN1A^-^/CDKN2A^+^*and double positive cells were selected (**Fig. 5A**) and pathway analysis performed. Senotype unique pathways revealed enrichment of immune regulatory pathways for *CDKN1A^+^/CDKN2A^-^* and *CDKN1A^+^/CDKN2A^+^* senotypes (**Fig. 5B**), which correlated with increased islet immune infiltrates (**Fig. 5C**), underscoring the pathophysiological relevance of our findings. Interestingly, the *CDKN1A^-^/CDKN2A^+^* SASP was enriched for cell signaling, differentiation and extracellular environment pathways and lower immune infiltrates, all in accordance with an adaptive identity.

**Figure 5.**
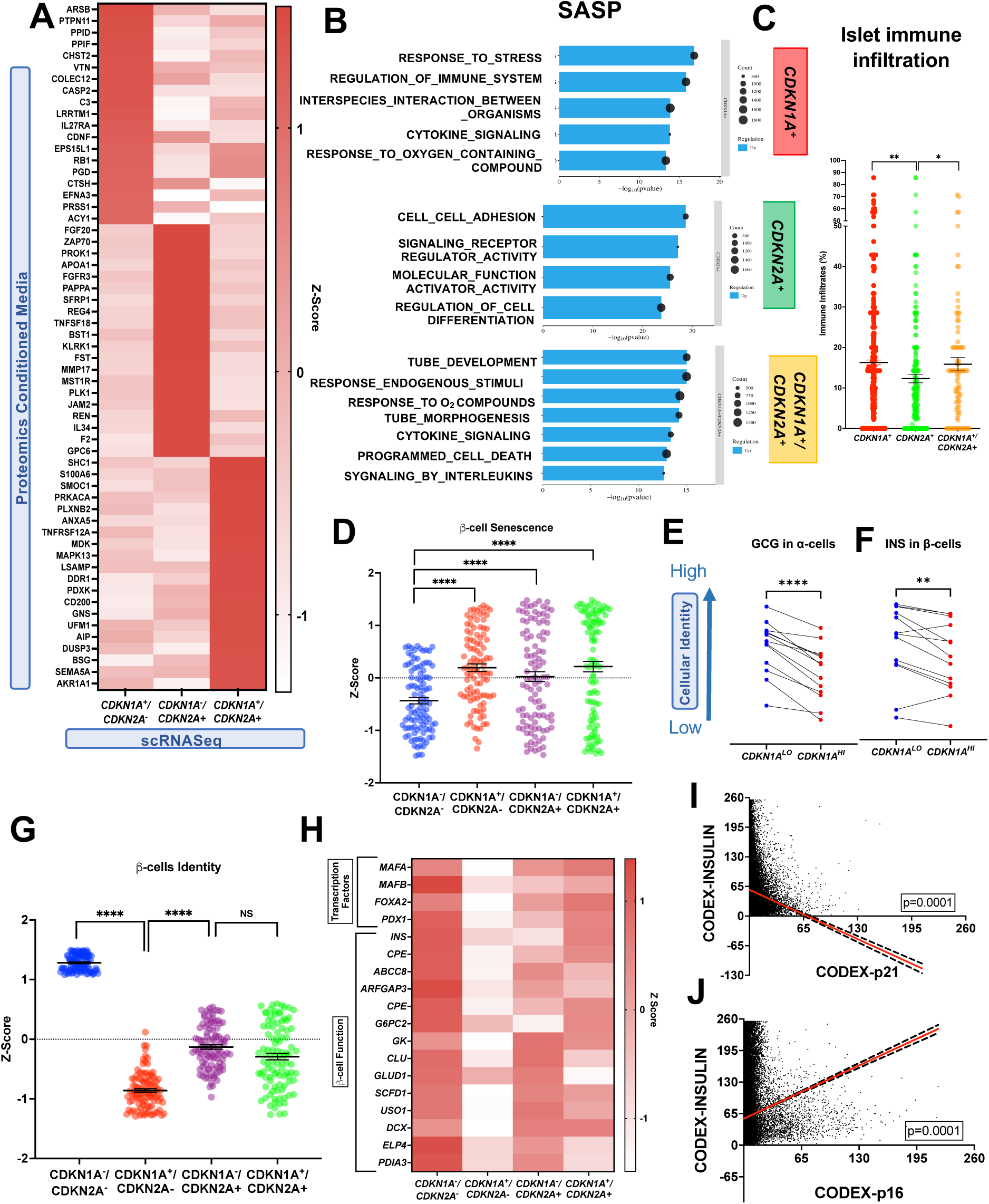
CDKN1A/p21 senotype specific proinflammatory SASP and cell identity changes. (**A**) Identification of senotype-specific SASP factors by integrating proteomic analysis of conditioned media from human islets and scRNASeq profile of specific senotypes in human β-cells. (**B**) Pathway analysis of senotype-specific SASP factors. (**C**) Immune infiltrates from integrated Xenium/CODEX datasets in islets with a specific senotype (**D**) Gene expression Z-score scatter plot of senescence genes in β-cell senotypes;. (**E, F**) scRNASeq data per donor GCG and INS expression across levels of *CDKN1A*, in α and β respectively. High and low expression of *CDKN1A*; *CDKN1A*^HI^ and *CDKN1A*^LO^. N=13, paired t-test, **p<0.01, ****p<0.0001. (**G**) Identity β-cell gene expression Z-score across senotypes. Mean+/-SEM, ****p<0.0001, n=13. (**H**) Heatmap of selected key β-cell identity and functional genes across senotypes. (**I**) Negative linear regression between p21 and INSULIN protein levels from β-cells as captured CODEX platform. Readings from all donors were pooled. (**J**) Positive linear regression between p16 and INSULIN protein levels from β-cells as captured CODEX platform. Reading from all donors were pooled.

### Senotype-specific changes in cellular identity

The effects of senotypes on the transcriptional identity and function of islet cells remain elusive.

A β-cell senescence score (**Suppl. Table 2**) was calculated for each senotype and double positive cells (**Fig. 5D**). All SnCs had comparable scores amongst each other which were significantly higher than in non-SnC cells. The development of a general non-cell type specific SASP score (**Suppl. Table 3**) also showed increased values in all three senotypes when compared with non-senescent β-cells (**Suppl. Fig. 6C**). These results confirm the senescent status of *CDKN1A^+/^ CDKN2A^-^*, *CDKN1A^-^ /CDKN2A^+^* and *CDKN1A^+^/CDKN2A^+^* β-cells.

Donor matched analysis of islet cells with different expression levels of *CDKN1A*, revealed loss of identity gene expression of *GCG* in α-cells, *INS* in β-cells **(Fig. 5 E, F)**, and *SST* δ-cells, but not γ-cells(**Suppl. Fig. A, B**) with development of SnC. Given that both increased and decreased expression of hallmark genes have been reported in senescent β-cells (SnCs)[3, 4], a deeper analysis of the transcriptional identity of this cell type is warranted. To evaluate the effects of SnC on cellular transcriptional identity, a β-cell specific hallmark and functional score was calculated (**Fig. 5G**) for all senotypes that revealed overall loss of cell identity. Nonetheless, the *CDKN1A^+/^ CDKN2A^-^* senotype had significantly lower identity score than CDKN1A-/CDKN2A+ β-cells. This was further highlighted by the dynamics followed by key β-cell transcription factors (*MAFA, PDX1*) and functional genes (*GK, ABCC8, CPE*) (**Fig. 5H**) whose expression was significantly lower than in *CDKN1A^-^ /CDKN2A^+^* and *CDKN1A^+^/CDKN2A^+^* β-cells. At the protein level, β-cell CODEX data was pooled for all donors and revealed a significantly negative correlation between p21 protein content and insulin protein levels (**Fig. 5I**), consistent with identity loss. In contrast, p16 protein β-cell levels, positively correlated with insulin (**Fig. 5J**), in line with previous publications.

P21-associated loss of identity in β and α cells in human pancreatic islets was reinforced using multiplex iCLAP images and quantification from pancreas of 4 younger donors and 3 older ones. Islet cells a was based on insulin and glucagon intensity from all samples (70 ROIs). Cells were classified into insulin+ β cells and glucagon+ α cells (**Fig. 6A**). Quantification of MAFA in p21+ and p21- β-cells, revealed significantly lower levels in the presence of the cell cycle inhibitor, irrespective of donor age (**Fig. 6B, D, E**). Similarly, p21+ α-cells had lower levels of identity transcription factor ARX in both age groups (**Fig. 6C, F, G**). Whereas the number of p21+ cells increased from 5.4% β-cells and 2.6% in α-cells in the younger donors (**Fig. 6H**) to 22% in β-cells and 17% in older ones (**Fig. 6I**), the identity loss with p21 expression was maintained across the lifespan. Representative pictures of 21-year old donor (**Fig. 6J-L**) revealed the relationship between the senescence marker p21 and loss endocrine cell identity and function. Whole-section overview (**Fig. 6J**) showed the distribution of insulin (β-cells), glucagon (α-cells), MAFA (β-cell transcription factor), ARX (α-cell transcription factor), p21, p16, and 53BP1. P21+/INS+ and p21+/GCG+ had diminished MAFA (β-cell identity and function) and ARX (α-cell identity) (**Fig. 6K,L**). Images from an 82-year old donor (**Fig. 6M-O**) revealed a higher occurrence of p21⁺/MAFA⁻ and p21⁺/ARX⁻ cells. These results show that p21 SnCs cells lose identity in islets irrespective of the donor’s age, a phenomenon with potential functional implications.

**Figure 6.**
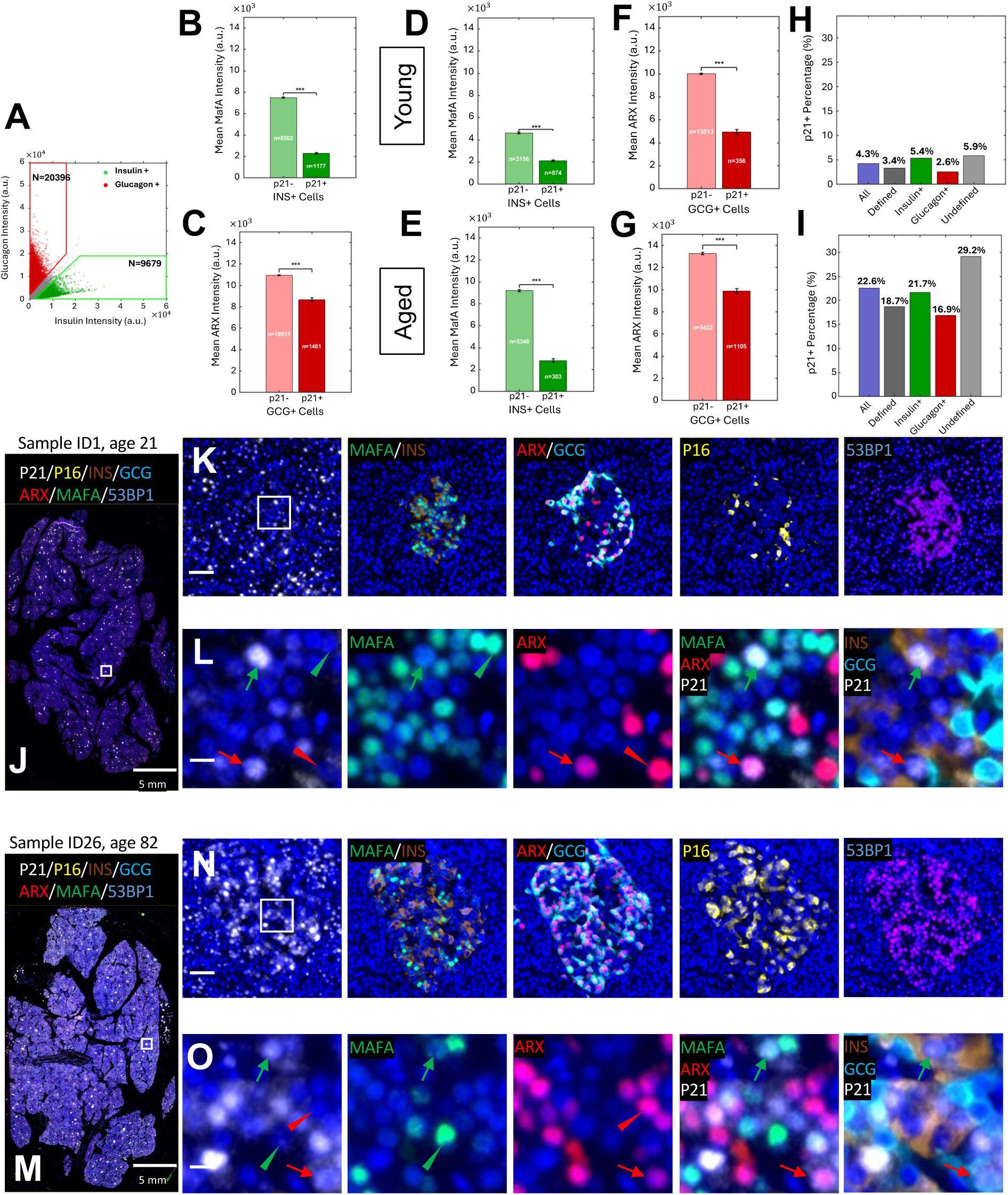
Senotype-specific loss of identity in β-cells and α-cells in human pancreatic islets across age. (**A**). Scatter plot showing islet cell classification based on insulin and glucagon intensity from all samples (70 ROIs, including 4 young and 3 old samples). Cells were classified into insulin+ β cells (green) and glucagon+ α cells (red), with undefined cells excluded from subtype-specific marker analysis. (**B. C**) Pooled analysis of total cells from all samples. Bar graphs show mean ARX intensity in glucagon+ α cells (C) and mean MafA intensity in insulin+ β cells (B) comparing p21− (p21 intensity < 20,000) and p21+ (p21 intensity > 20,000) cells. **D, F**. Analysis of the 4 young samples (40 ROIs). Defined cells are sum of insulin+ cells and glucagon+ cells. show mean MafA intensity in insulin+ β cells, ARX intensity in glucagon+ α cells and respectively, comparing p21− and p21+ cells. **E,G**. Analysis of the 3 old samples (30 ROIs) using the same measurements as in **D, F**. Across both young and old samples, p21+ β cells showed reduced MafA intensity and p21+ α cells showed reduced ARX intensity compared with p21− cells, suggesting that increased p21 expression is associated with loss of islet cell identity marker expression. ***p < 0.001**. H** shows the percentage of p21+ cells across all cell types in young and older (I) donors. p21+ cells were defined as cells with p21 intensity >15,000 a.u., while p21− cells were defined as cells with p21 intensity <15,000 a.u. (**J-L)** Multiplex iCLAP images from a young donor (age 21) illustrate the relationship between the senescence marker p21 and endocrine cell identity and function. Whole-section overview (**J**) shows the distribution of insulin (β-cells), glucagon (α-cells), MAFA (β-cell transcription factor), ARX (α-cell transcription factor), p21, p16, and 53BP1. Higher-magnification panels (**K**) display single-senescence marker maps for p21, p16 and 53BP1, together with colocalized INS/MAFA and GCG/ARX. (**L**) Single-cell insets highlight representative p21^+^/INS^+^ (green) and p21^+^/GCG^+^ (red) cells (arrow) with diminished MAFA (β-cell identity) and ARX (α-cell identity). (**M-O**) Multiplex iCLAP images from an old donor (ID26, age 82) illustrate the relationship between the senescence marker p21 and endocrine cell identity and function. Whole-section overview (**M**) shows the distribution of insulin (β-cells), glucagon (α-cells), MAFA (β-cell transcription factor), ARX (α-cell transcription factor), p21, p16, and 53BP1. Higher-magnification panels (**N**) display single-senescence marker maps for p21, p16 and 53BP1, together with colocalized INS/MAFA and GCG/ARX. Single-cell insets (**O**) highlight representative p21^+^/INS^+^ (green) and p21^+^/GCG^+^ (red) cells (arrow) with diminished MAFA (β-cell identity) and ARX (α-cell identity). High occurrence of p21⁺/MAFA⁻ and p21⁺/ARX⁻ cells are found in islet derived from older donor.

Identity analysis supported the existence of a *CDKN1A^+^/*p21 senotype characterized by loss of identity and a maladaptive profile of senescence, and a *CDKN2A^+^*/p16 senotype which conserved transcriptional identity. As a reference, senescence and SASP transcriptional signatures were developed for all endocrine cell types (**Suppl. Tables 2 and 3**).

### *CDKN1A^+/^ CDKN2A^-^* β-cells were dysfunctional as evaluated by insulin secretion

The function of isolated pancreatic islets was evaluated after overnight recovery with static and dynamic glucose stimulated insulin secretion (GSIS) (**Fig. 7A**) and donor-matched to scRNASeq results for *CDKN1A^+^/CDKN2A^-^* and *CDKN1A^-^/CDKN2A^+^* transcript levels in β-cells. Islet preparations were grouped based on *CDKN1A* levels in β-cells (**Fig. 7B**) with no significant differences in *CDKN2A* levels between groups. Static GSIS revealed that β-cells with low levels of *CDKN1A* (*CDKN1A*^LO^) were functional and had increased insulin secretion at high glucose concentrations, whereas those with higher levels of CDKN1A (*CDKN1A*^HI^) were dysfunctional (**Fig. 7C**) and had a lower secretion index (**Fig. 7D**). Additional assessment of β-cell function was performed with dynamic GSIS by islet perifusion. Islets were divided into two groups based on *CDKN1A* levels in β-cells and with no significant differences in *CDKN2A* levels (**Fig. 7E**). Islet perifusion confirmed dysfunction of *CDKN1A*^HI^ cells with abolished responses to both high glucose levels (16.8 mM glucose) and to the secretion potentiator, 3-isobutyl-1-methylxanthine (IBMX) (**Fig. 7F**).

**Fig. 7.**
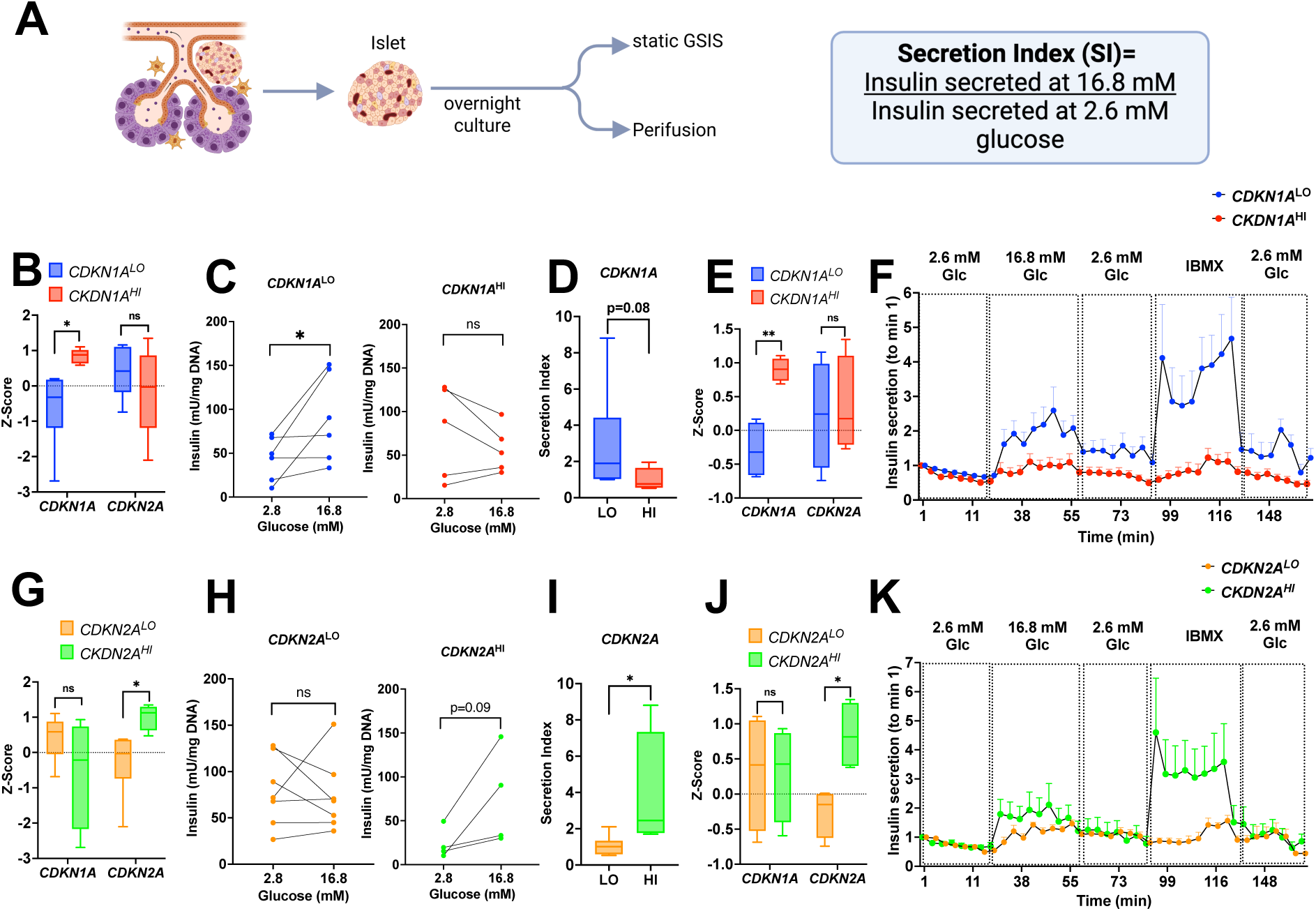
Loss of function in *CDKN1A* senotype leads to impaired insulin secretion from human islets. (**A**) Graphical abstract of functional evaluation of isolated human islets; (**B**) *CDKN1A* and *CDKN2A* gene expression levels in islets preparation used for functional evaluation with GSIS with low (LO n=6) and high (HI n=5) levels of *CDKN1A* as assessed by scRNASeq. Mean+/-SEM, *p<0.05. (**C**) Static GSIS in *CDKN1A*^LO^ and *CDKN1A*^HI^ (**D**) Secretion Index in *CDKN1A*^LO^ and *CDKN1A*^HI^. t-test, *p<0.05. (**E**) *CDKN1A* and *CDKN2A* gene expression levels in islets preparation with low (LO n=4) and high (HI n=4) levels of *CDKN1A* as assessed by scRNASeq used for functional evaluation with dynamic islet perifusion. (**F**) Dynamic insulin secretion normalized by insulin levels at min 1 of *CDKN1A*^LO^ and *CDKN1A*^HI^ islet preparations. (**G**) *CDKN1A* and *CDKN2A* gene expression levels in islet preparations with low (LO n=7) and high (HI n=4) levels of *CDKN2A* as assessed by scRNASeq used for functional evaluation with GSIS. Mean+/-SEM, *p<0.05. (**H**) Static GSIS in *CDKN2A*^LO^ and *CDKN2A*^HI^; (**I**) Secretion Index in *CDKN2A*^LO^ and *CDKN2A*^HI^. Mean+/-SEM, *p<0.05. (**J**) *CDKN1A* and *CDKN2A* gene expression levels in islets preparation with low (LO n=4) and high (HI n=4) levels of *CDKN2A* as assessed by scRNASeq used for functional evaluation with dynamic islet perifusion (**K**) Insulin secretion normalized by insulin levels at baseline (min 1) of *CDKN2A*^LO^ and *CDKN2A*^HI^ islet preparations.

To assess the effects of *CDKN2A* in β-cell function, islets were sorted based on transcript levels into a *CDKN2A*^LO^ and a *CDKN2A*^HI^ group (**Fig. 7G**). In accordance with previous reports [3] *CDKN2A*^HI^ islets had a better secretory response (**Fig. 7H**) and a higher secretion index (**Fig. 7I**), both indicating improved function. Perifusion studies confirmed the functionality of *CDKN2A*^HI^ islets (**Fig. 7J, K**)

## DISCUSSION

The comprehensive multi-omic and functional analysis of the human endocrine pancreas revealed that the most significant changes during the human lifespan are a decrease in the proportion of endocrine β-cells, insulin, and C-peptide coupled with an increase in *CDKN1A* and *CDKN2A* pointing to β-cell senescence as a hallmark of human aging.

The discrete distributions of *CDKN1A* and *CDKN2A* in spatial proteomic and transcriptomic analyses at single cell resolution led to the identification of two senotypes. The *CDKN1A^-^/CDKN2A^+^* β-cell senotype was characterized by maintenance of transcriptional identity, a low inflammatory SASP, and preserved function and its dynamics can be followed with HMGB1 as a biomarker. In contrast, *CDKN1A^+^/CDKN2A^-^* cells were characterized by downregulation of hallmark and functional β-cell genes, dysfunction as seen by lack of GSIS, and an immune regulatory SASP, which associated with increased islet immune infiltrates. Spatially, *CDKN1A^+^/CDKN2A^-^* β-cells associated with acinar cells and were closer to endothelial cells, suggesting important tissue cross talk between senotypes. Interestingly, double positive *CDKN1A^+^/CDKN2A^+^*β-cells had mixed characteristics with maintenance of transcriptional identity and an immune regulatory SASP.

Based on these attributes, we suggest the existence of an adaptive senotype comprising of *CDKN2A*^+^ β-cells that maintains terminal differentiation fate and can modulate glycemia through appropriate insulin secretion in response to stimuli, with minimal deleterious effects on neighboring cells. This is consistent with previous reports showing that p16 overexpression in β-cells improved expression of terminal differentiation genes (*MAFA, GK, PDX1*) and insulin secretion [3].

In contrast, *CDKN1A^+^* β-cells represent a maladaptive senotype characterized by loss of transcriptional cell identity and dysfunction. The metabolic implications of this phenomena are important and they are induced by age and insulin resistant states, such as a high BMI. Relevant effects of this senotype on neighboring tissues are underscored by high *CDKN1A* levels in the acinar-islet border, consistent with studies showing that activation of the insulin receptor (INSR) induces senescence in various cell types [14–16].

Increased proximity of *CDKN1A*^+^ β-cells to extra-islet endothelial cells is consistent with reports of SnC endothelial cells inducing senescence and metabolic dysfunction in metabolically relevant tissues [17]. A bidirectional SASP cross talk between endothelial and β-cells hypothesizes endothelial SASP induction of β-cell SnC through the p21 pathway and the SASP from *CDKN1A^+^/CDKN2A^-^* β-cells promotion of islet immune infiltration from the intravascular space.

The senotype-specific SASP at the transcriptional level included factors previously described in other tissues. For example, PAPPA was detected as part of the *CDKN1A-/ CDKN2A*+ SASP. PAPPA is a known longevity regulator that impacts the IGF1 signaling pathway [18]. This SASP profiling, along with islet-cell specific senescence and SASP transcriptional signatures (**Suppl. Tables 2 and 3**), will provide resources to further identify commonalities and differences among endocrine pancreatic senotypes.

Changes in cell composition of SnCs in islets deserves consideration due to its potential effects on β-cell function. We found a significant age-dependent decrease of β-cells and δ-cells in senescent islets; α-cells were not downregulated with age while there was an increase in γ-cells. A decrease in δ-cells predicts lower intra-islet somatostatin which negatively modulates glucagon and insulin secretion [19]. Lower inhibitory signals could lead to increased glucagon secretion from α-cells reflected as the hyperglucagonemia observed in individuals with Type 2 diabetes. This pathophysiological change can be explained by two factors: the loss of insulin’s inhibitory effect on α-cell secretion and β-cell dysfunction [20]. The increase of γ-cells in SnC islets is intriguing given that this cell type is preferentially located in the inferior head of the pancreas due to its distinct embryological origin. However, the systematic dissection of the pancreas and omics profiling of multiple samples per donor representing different pancreatic sections should offset a potential a sampling bias. Instead, given the loss of identify in SnCs, increased detection of PP could be due to off-target expression of this gene.

Islet structural changes with age can also have functional repercussions. Pancreatic islets have a portal circulatory system where flow has been shown to go from the central β-cell core to the periphery, thereby signaling α, δ, and γ- cells [21]. Higher circularity and lower eccentricity, as was seen in donors under 45 years of age, would predict an even exposure of islet cells to insulin thereby providing feedback signals. However, with age eccentricity increased. This predicts there may be altered exposure of regulatory endocrine cells to insulin secretion.

Together, these findings define distinct and functionally divergent senotypes within the human pancreas, revealing that senescence is not a uniform cellular state but a heterogeneous program with opposing consequences for islet biology. We identify a maladaptive *CDKN1A*⁺ senotype characterized by loss of cellular identity, proinflammatory SASP signaling, and impaired endocrine function, in contrast to a more adaptive *CDKN2A*⁺ state associated with preserved cellular integrity. These results establish a mechanistic framework linking senescence heterogeneity to pancreatic dysfunction during aging and metabolic disease, while providing a foundation for precision senotherapeutic strategies aimed at selectively targeting pathogenic senescent cell populations in type 2 diabetes.

### Limitations

The following limitations are acknowledged:

1. This study was performed using solely high-quality human pancreas with single-cell resolution at the transcript and protein level. However, some reported phenotypes remain to be accounted for. For example, expression of p21 in β-cells was shown to be protective in models of type 1 diabetes [5], while the endothelial p16 population has been shown to be deleterious to neighboring cells through its SASP [17]. We believe some of these discrepancies are due to the nature of the stressor which induces β-cell senescence, a topic out of the scope of the current manuscript.
2. The cross-sectional nature of this study cannot address previously described temporal relationships between *CDKN1A* and *CDKN2A* populations [22].
3. The size limitation in the cohort of donors is counterbalanced by the deep sequencing of multiple pancreatic regions per specimen. Additionally, the personal omics approach ensures that every organ has many data points from orthogonal techniques providing a deep understanding of senescence biology.
4. The role and changes that these senotypes undergo in metabolic diseases such as Type 2 Diabetes, remain to be determined. Based on this blueprint, *CDKN1A*^+^ β-cells are predicted to be increased and actively involved in the development of the disease in humans.

Further studies and integration of databases from different organs will address some of these unresolved questions. SenNet has generated the necessary resources and tools for the development of the next generation of SnC biology and further elucidation of the relevance of SnCs in human health and disease.

## ACKNOWLEDGMENTS

First and foremost, we are grateful to all the donors and their families. The authors thank Angela Wood from the Flow Cytometry Core.

Dr. Susan Bonner-Weir and Dr. C. Ronald Kahn for crucial input and discussions. Purified islets, ducts and acinar tissue were provided by Prodo Laboratories, *Inc*., in Aliso Viejo, CA.

## FUNDING

This work was funded by the NIH/NIA U54AG075941 to the SenNet Initiative, KAPPSen Tissue Mapping Center. This work also received support from National Institutes of Health (NIH) grants 1R01DK132535 to CAM, P30 DK036836 to Joslin Diabetes Center (Cores), UH3CA275681 to PHW, U54 AG079758 to PA, R37AG013925 to JLK, R33AG061456 to JLK, R33AG061456 to JLK, Thomas J Beatson Jr Foundation grant 2025-020 and the Richard and Susan Smith Family Foundation Award to CAM, and Hevolution Foundation grant HF-GRO-23-1199148-3, the Connor Fund, and Robert J. and Theresa W. Ryan to JLK.

## AUTHOR CONTRIBUTIONS

K.I, H.P, J.D, S.D, D.B, J.A.D, F.W., P.R., G.K., N.M., J.L.K., T.T. and C.A.M. conceived the project and wrote the manuscript; K.I, H.P, J.D, S.D, D.B, P.C, C.C, S.L, F.H, J.A.D, G.N.E, F.W, B.Y, P-H.W, and C.A.M. researched data. P.C, J.L.W, P.A, D.U, J.L.K, and G.A.K provided critical discussions and edited the manuscript. D.W, S.P, A.P, G.A, J.H.C, V.D.G, T.T, N.M, and P.R provided critical discussion. All authors reviewed the manuscript.

## COMPETING INTEREST DECLARATION

The authors declare no financial/non-financial competing interest.

## DATA AVAILABILITY

Datasets analyzed in this study are publicly available through the SenNet Data Sharing Portal (https://data.sennetconsortium.org/) under open or controlled access via dbGaP. Three collections were used.

**SNT793.SZRS.468** provides multimodal pancreatic tissue data, including 32 Xenium transcriptomics datasets, 28 PhenoCycler images, and 32 H&E sections.

**SNT566.NMTV.379** contains 30 single-cell RNA-seq datasets generated with the 10x Genomics Chromium platform.

**SNT947.NXPB.793** 10x Visium and H&E stain profiling of pancreas.

## Notes

### Competing Interest Statement

The authors have declared no competing interest.

