## Supplementary material for "Distinct senescent β-cell senotypes differentially drive islet aging and dysfunction": Online Methods

**Whole Pancreas**

**Ethics statement and sources**

For whole pancreas organ was collected from UT Health San Antonio (UTHSCSA), the Center for Life Donor Biorepository was determined to be Non-Human Research by the UTHSA IRB, as the donors are deceased at time of inclusion. The pancreata were obtained through this biobank protocol at University Hospital Center for Life, which is managed in partnership between University Health Transplant Institute (UHTI) and Texas Organ Sharing Alliance (TOSA), who is the local Organ Procurement Organization. TOSA has jurisdiction over the donors, and an approval was received from their Advisory Board to receive organs intended for research.

Donor pancreatic organs were procured by the Center for Organ Recovery & Education (CORE), Pittsburgh, under appropriate research consent, and processed by Imagine Islet Center (Pittsburgh) in accordance with institutional and regulatory guidelines.

For dispersed pancreas, removal of the tissue specimens from the donors and their use for research was approved by PRODO Laboratories, Inc. Informed consent was obtained through the DMV donor registry and/or next of kin.

Research conducted using tissues obtained through these programs, was determined to meet the characteristics of Not Human Subjects Research (NHSR).

**Organ Procurement and Specimen Handling Procedures**

The Texas Organ Sharing Alliance (TOSA) followed standardized recovery instructions to ensure procurement of an intact pancreas with all necessary anatomical landmarks preserved for research orientation. A portal and arterial system flush with either Histidine–Tryptophan–Ketoglutarate (HTK) or University of Wisconsin (UW) preservation solution was performed *in situ* whenever possible. The duodenum was stapled and the common bile duct lumen was marked with a long silk suture, and the portal vein lumen is marked with a long Prolene (or equivalent) suture to facilitate subsequent anatomical identification. Priority was given for pancreatic parenchyma to remain fully intact.

Following recovery, the pancreas was packaged in accordance with standard clinical transplant procedures using Organ Procurement and Transplantation Network (OPTN)–supplied disposable shipping containers. Each container consists of an outer corrugated plastic or corrugated cardboard shell with a minimum burst strength of 200 lb and coated with a water-resistant material, a water-tight and secured opaque plastic liner positioned between the outer and inner containers, and an inner insulated container with a 1.5-inch wall thickness (or equivalent thermal resistance) containing sufficient cooling material to maintain appropriate temperature during transport. Within this insulated compartment, a sealed plastic liner encloses the cooling medium; for this study, ice was used. The pancreas was immersed in HTK or UW preservation solution and sealed in a sterile polyethylene bag, which was then placed into a rigid polyethylene container. This container was subsequently positioned inside the internal sealed liner and fully surrounded by cooling material to ensure optimal preservation conditions during transit.

**Specimen Receipt and De-identification**

Specimens recovered through Center for Life (CFL) procurements are packaged on-site and transported directly to the University of Texas Health Science Center at San Antonio (UTHSCSA) CFL Biorepository, located at University Hospital (UH). Specimens recovered through the Texas Organ Sharing Alliance (TOSA) were transported to the biorepository by the Nationwide Organ Recovery Transport Alliance (NORA). Upon receipt, transplant research personnel remove all UNOS identifiers and any associated personal health information. A CFL study-specific identification number was then assigned prior to specimen allocation. The de-identified specimen was shipped by FedEx Priority Overnight. Research conducted using tissues obtained through this program from brain-dead donors has been determined to be Not Human Subjects Research (NHSR).

**Inclusion and Exclusion Criteria**

In both whole pancreas and dispersed pancreas donors from age over 20 in both genders, all ethnicities were included. Donors with these were excluded; DM, HTN with end-organ damage, BMI>30, GFR<90 ml/min/1.73 m^2^, cardiovascular disease, pancreatitis, malignancy, chronic pulmonary, hepatic, infectious, gastrointestinal, endocrine, neurologic, or inflammatory disease. Also chronic use of systemic steroids or use of agents with senolytic activity was excluded. As it was difficult to recruit donors, Two donors with BMI over 30 were accepted; 30.2 and 30.9 in dispersed pancreas experiment

**Specimen sectioning**

The whole pancreases from heart beating brain dead donors were procured by the University of Texas Health San Antonio or Imagine Pharma, and shipped to Joslin Diabetes Center, University of Harvard Medical. The pancreases were cut into the Head (Superior and Inferior), Body, and Tail of the locations following the guidelines published in the following surgical guidelines [1], cut into 4 mm thickness slices, and placed into paraffin cassettes. The paraffin cassettes were fixed in 4% paraformaldehyde for 24 hours and processed for paraffin embedding at Joslin or BIDMC Histology Core. At least three paraffin cassettes per each location of Superior Head, Inferior Head, Body, and Tail were shipped to The Jackson Laboratory; all the procedures were done at 4°C and exposure to direct light was avoided.

**FFPE Hematoxylin and Eosin (H&E) Staining**

H&E staining was performed following a published protocol (dx.doi.org/10.17504/protocols.io.6qpvr3653vmk/v1). FFPE blocks were stored at 4 °C and sectioned under RNase-free conditions for spatial transcriptomics. After sequential incubations in *xylene*, graded *ethanol*, and staining steps, slides were scanned using a *NanoZoomer HT2.0* (Hamamatsu) at 40× brightfield. Images were analyzed using *NDP software*.

**FFPE Block DV200 Evaluation**

RNA quality from FFPE tissue was assessed prior to use following the Visium Spatial Gene Expression for FFPE protocol. Two to four 5 µm sections were collected under RNase-free conditions, and RNA was extracted using *RNeasy Micro Kit* (Qiagen, Cat. No. 74004). RNA integrity was evaluated using *Tapestation High Sensitivity RNA ScreenTape* (Agilent), and DV200 scores were calculated as the percentage of RNA fragments >200 bases using *TapeStation Analysis software*. All blocks achieved DV200 >50%.

**Visium Spatial Transcriptomics with CytAssist**

Sections (6.5 mm × 6.5 mm) containing regions of interest were mounted on *ColorFrost Plus slides* (Fisher), deparaffinized, H&E stained, and imaged using a *NanoZoomer SQ* (Hamamatsu). Slides were hybridized with human-specific probe sets (10x Genomics), and probes were transferred to a *Visium CytAssist Slide* using the *CytAssist* instrument (10x Genomics), followed by library preparation per manufacturer’s protocol (CG000495).

Libraries were quantified using *Tapestation High Sensitivity DNA ScreenTape* (Agilent), fluorometry (*Qubit*, ThermoFisher) and verified by *KAPA qPCR*. Sequencing was performed on an *Illumina NovaSeq X+* 10B flow cell (100 cycles) using a 28-10-10-90 read configuration, targeting 100,000 read pairs per tissue-covered spot. Base call files were converted to FASTQ using *bcl2fastq* v2.20.0.422 (Illumina).

Brightfield and CytAssist images were aligned manually using *Loupe Browser* v6.4.1 via landmark registration. Whole-slide images were uploaded to a local *OMERO* server, and rectangular regions of interest (ROIs) were defined using *OMERO.web*. OMETIFF images of each ROI were generated programmatically using the *OMERO Python API*. FASTQ files, image registration JSON, and OMETIFF images were processed using *Space Ranger count pipeline* v2.1.0 (10x Genomics) with the GRCh38-specific filtered probe set (10x Genomics Probeset v1.0.0).

**Multimodal Imaging Assays (Xenium + Phenocycler-Fusion + H&E)**

FFPE tissue slides were first processed for *Xenium In Situ Gene Expression* (10x Genomics) following manufacturer protocols (CG000580, CG000582, CG000584). Briefly, sections underwent deparaffinization, decrosslinking at 80 °C, probe hybridization, ligation, amplification, and autofluorescence quenching before imaging on the *Xenium Analyzer*.

Slides were either processed immediately or stored for up to 2 weeks at 4 °C in 50% glycerol/PBS. For *Phenocycler-Fusion* (Akoya Biosciences), slides were washed in PBS and Milli-Q water, followed by antigen retrieval in citrate (pH 6.0) or Tris-EDTA (pH 9.0) buffer using a pressure cooker (90–110 °C, 15 min). After cooling and rinsing, multiplexed antibody staining was performed per the Phenocycler-Fusion User Guide (PD-000011 Rev L).

Following multiplexed imaging, slides were processed for H&E staining. Flow cells were removed by incubating slides in *xylene* or *Histo-Clear* for 24 h at room temperature. Slides were rehydrated through graded ethanol (100%, 95%) and rinsed in water. Staining was performed sequentially with *Mayer’s Hematoxylin* (4 min), *Bluing Reagent* (1 min), and *Alcoholic Eosin* (2 min), followed by dehydration in ethanol and clearing in xylene. Slides were mounted using *DPX mountant* and cured for at least 20 min before imaging.

**Islet composition analysis**

To find islets in all tissues we started with Xenium data where each cell is annotated with a cell type, *e.g*., endocrine-beta cell. We built a KDTree with cell centroids spatial location x and y for fast search of cells neighborhoods. For each endocrine cell we found its neighboring endocrine cells within radius of 16 um (maximum allowed center to center distance). Then we constructed the neighborhood graph and partitioned it into components with at least 10 cells. For each component we added non-endocrine neighboring cells (within 16 um distance) except acinar cells. We computed each islet morphological and molecular properties: area, perimeter length, eccentricity, density, centralization, localization, mean nucleus area, mean cell area, mean number of transcripts per cell, proportion of each cell type, average expression of each gene, and each CODEX channel across cells.

To compute cell type centralization for a given cell type, we take cells belonging to an islet with spatial $x$ and $y$ coordinates denoted as $\left\{ c_{i}, \right\}$, we compute its centroid $\bar{c}_{\text{cell}\text{type}}=\frac{1}{n}\sum_{i} c_{i}$; and the centroid of the islet $\bar{c}_{\text{islet}}$ over all cells in that islet. Centralization quantifies how close the cell-type centroid is to the islet center, normalized by the size of the islet. Specifically, we measure the Euclidean distance $d=\parallel\bar{c}_{\text{cell}\text{type}}-\bar{c}_{\text{islet}}\parallel$ and divide by the 99th percentile of all distances from $\bar{c}_{\text{islet}}$ to cells in the islet, $q_{0.99}$, to obtain a robust, outlier-resistant normalization. The final score is

$$C=1-d/q_{0.99}$$

with $C=1$ indicating perfect central alignment of a cell type within an islet and $C=0$ indicating a cell type centroid at the islet’s periphery.

To obtain cell type localization, we compare the within-group spatial distribution of the cell type to that of the encompassing islet. Specifically, we compute the mean pairwise Euclidean distance among all points in the component, $a$, and among the cell type points, $b$. To ensure the metric remains bounded and interpretable when the subgroup is more dispersed than the component, we cap $b$ at $a$ (i.e., if $b>a$, set $b=a$). The localization score is then

$$L=(a-b)/(a+b),$$

yielding values in $\left[ 0, 1 \right]$, where larger values indicate tighter grouping of the cell type relative to all the component cells with $L=0$ meaning equal distribution. If either set has fewer than two points, $L$ is set to 0.

For each tissue we computed trimmed medians using global 0.1%/99.9% quantile bounds of each variable. Then we computed Pearson correlation coefficient between all pairs of variables. The hierarchical clustering within each heatmap group of variables was done with Euclidean distance and Ward linkage method. The variables were separated into categories such as demographic, cell type morphology, CODEX protein, and Xenium genes.

To find p16 or p21-high islets, we used a cutoff of 2.0 on the normalized fluorescence intensity of the CODEX channels. We separated subsets of islets to a particular age group, *e.g*., young, and hierarchically clustered islets according to their endocrine cell type proportions. The clustering metric was Pearson correlation and average linkage method. The clustering threshold of 0.25 was used to differentiate groups of islets with different endocrine cell type compositions. *CDKN2A* probes in Xenium and Visium recognized both the INK and ARF transcripts.

**iCLAP protein detection**

Johns Hopkins University received ten slides per sample for each of the eight tissue specimens (Donors 1, 3, 5, 7, 22, 24, 26, and 27) collected by Harvard University. Four-micron paraffin sections were baked at 42 °C for 3 hours and dried overnight at room temperature in a desiccator. Slides were then dewaxed in xylene, rehydrated through graded ethanol, and finally rinsed several times in water. For antigen retrieval, slides were placed in a heat-resistant plastic container filled with antigen retrieval buffer (Vector Laboratories, H-3300-250) and heated for 20 minutes using a food steamer (Bella). Tissue sections were multiplex-labeled using antibodies against p16 (Roche Diagnostics, 705-4793; ready-to-use), p21 (BD Biosciences, 554228; dilution 1:500), ARX (Abcam, ab308260; dilution 1:500), MAFA (Invitrogen, MA5-44325; dilution 1:200), 53BP1(Bethyl Laboratories, A700-011; dilution 1:500), insulin(Abcam, ab309368; dilution 1:250), and glucagon(Abcam, ab313316; dilution 1:250). Insulin and glucagon were detected using fluorophore-conjugated antibodies, while the remaining markers were detected using tyramide signal amplification (TSA). Staining and imaging were performed over three sequential rounds to detect all seven markers. In the first round, p16 and p21 were stained together with Hoechst 33342 for nuclear counterstaining. In the second round, ARX and MAFA were stained. In the third round, insulin, glucagon, and 53BP1 were stained. Between each staining round, fluorescent signals were quenched by placing the slides in a transparent container containing bleaching solution (2 M H₂O₂ and 3 mM EDTA in PBS, adjusted to pH 12.5) and positioning the container between two 5000-lux LED light pads (HSK, 615517997868) for 1 hour. After each round, stained slides were imaged using an inverted fluorescence microscope (see Fluorescence Microscopy section for details).

**Fluorescence microscopy**

Fluorescently labeled tissue sections were imaged using a Hamamatsu Flash 4.0 CMOS camera mounted on a Nikon Ti-E inverted research microscope. The microscope was equipped with a motorized stage and motorized excitation and emission filter wheels, all controlled through NIS-Elements (Nikon). A Lumencor SpectraX 6 light engine served as the illumination source. For each sample, a custom imaging grid was generated to capture the entire tissue area using an S Fluor 10× objective (NA 0.5; MRF00100, Nikon). Adjacent fields were acquired with a 10% overlap to enable accurate stitching. The Perfect Focus System (Nikon) was used throughout acquisition to maintain a stable focal plane across large tissue regions. Under this optical configuration, the acquired images had a pixel size of 0.65 μm. Images within each grid were stitched using previously described algorithms [2].

**Tissue image registration**

To align images acquired across different staining rounds, we applied the following registration workflow. Nuclear images (i.e., the DAPI channel) from each round were used as the reference for alignment. The registration procedure consisted of two main steps: an initial global rigid registration, followed by a local grid-based deformable registration, as previously described [3]. Global rigid alignment was performed on images downsampled by a factor of five to improve computational efficiency, whereas local deformable registration was carried out on the full-resolution data to maximize spatial accuracy. The deformable registration employed a grid spacing of 325 μm. Aligned whole-slide images were exported in OME-TIFF format using the libvips library [4] and visually inspected in QuPath [5] for quality assessment.

**Dispersed Pancreas (islet, acinar, duct cells)**

**Pancreas dispersion (PRODO)**

Upon arrival, the pancreas was removed from the transportation container and placed in a new stainless-steel dissection pan with 500 mls of sterile saline (Baxter, NDC 0338-0048-04) that was kept cold with ice packs (Uline, S-7361 (3 cold & 3 frozen) beneath the pan. The extra pancreatic fat, along with the spleen residual was surgically dissected off the pancreas. The duodenum was also removed and the organ was sterilized before moving to the distention phase. The pancreas distension step was accomplished manually with the pancreas placed into an empty stainless-steel pan and manually injected with a digestive enzyme. Once pre-digestion had occurred, the remaining fat was dissected off and the pancreas was transferred to the digestive chamber. During this phase, the digestive enzyme was reintroduced so the pancreas slowly digested and the cells, once free, traveled through a circuit to collection conicals, which were kept at 4C. The collection conicals were then spun down and the pellets combined for an incubation period.

After the incubation period, the cells were purified using a density gradient, along with a COBE machine. Purified islets floated to the top, while ductal cells were in the middle (acinar and islets are present) and acinar sank to the bottom. These layers were collected in thirty-three tubes, separating pure islets, mixed-purity duct cells, and purified acinar cells. The various cell types were then cultured in flasks and incubated until they had recovered for shipping. Equivalent amounts of dispersed islets, acinar cells, and ductal cells from healthy donors were shipped to Joslin Diabetes Center and Jackson Laboratory.

**Preparation of Single-Cell Suspensions**

Pancreatic acinar and ductal cells were isolated and transferred to *PIM(R)® medium* (Prodolab) supplemented with 10% *fetal bovine serum* (FBS; HyClone). After three washes in *phosphate-buffered saline* (PBS), tissues were digested with *TrypLE™ Express* (GIBCO) at 37 °C for 30 min (acinar) or 15 min (ducts), with gentle pipette dissociation every 5 min. Digestion was terminated using *Dulbecco’s Modified Eagle Medium* (DMEM; GIBCO) containing 10% FBS and 1:100 Glutamate, followed by centrifugation (1300 rpm, 4 °C, 2 min) and resuspension in stop medium.

Pancreatic islets were washed with PBS and digested using *StemPro™ Accutase™* (1 ml per 1,000 islets) for 10 min at 37 °C. The reaction was quenched with *CMRL 1066*, and cells were centrifuged (230 × g, 5 min) and resuspended in CMRL. All cell suspensions were filtered through 40 µm *Flowmi® strainers* (Sigma) and counted using AO/PI viability assay (*Luna-FL automated counter*).

Cells were fixed for single-cell RNA sequencing following 10x Genomics protocol CG000478. Briefly, single-cell suspensions were centrifuged (350 rcf, 4 °C, 5 min), resuspended in *Fixation Buffer*, incubated for 1 h at room temperature, and stored at 4 °C for 16–24 h. After centrifugation (850 rcf, 5 min, room temperature), *Quenching Buffer* was added on ice. Cell concentration was determined using AO/PI staining, and pre-warmed *Enhancer* (10x Genomics PN-2000482) was added. Samples were stored for up to one week before processing.

**Single-Cell Library Preparation and Sequencing**

After dissociation, cells were washed and resuspended in *PBS* containing 0.04% *BSA* and processed immediately. Cell viability was assessed using a *LUNA FX7 automated cell counter* (Logos Biosystems), and up to 12,000 cells per suspension were loaded onto one lane of a *10x Genomics Chromium Chip G*.

Single-cell capture, barcoding, and library preparation were performed on the *Chromium X platform* (10x Genomics) using NEXTGEM chemistry (v3.1) according to the manufacturer’s protocol (#CG000315). cDNA and libraries were quality-checked using *Tapestation 4200* (Agilent) and *Qubit Fluorometer* (ThermoFisher), quantified by *KAPA qPCR*, and sequenced on an *Illumina NovaSeq X+* 10B flow cell (100-cycle), with a 28-10-10-90 asymmetric read configuration. Libraries targeted ~6,000 barcoded cells at an average depth of 50,000 reads per cell.

Illumina base call files were converted to FASTQ using *bcl2fastq* v2.20.0.422 (Illumina). Gene expression FASTQs were aligned to the *GRCm38* reference genome with vM23 annotations (GENCODE) using the *Cell Ranger multi pipeline* v8.0.0 (10x Genomics) and the mm10 reference (2020-A).

**scRNASeq Cell identity evaluation**

First, GCG, INS, and SST were selected as indicators of cell identity for alpha, beta, and delta cells, respectively. Then, for each cell type, cells of each donor were split into five quintiles based on CDKN1A expression. The highest three quintiles were selected as p21-High, -Mid, and -Low categories. Donor-level mean indicator gene levels were compared across the categories using the paired t-test.

**FACS analysis with βGal**

After overnight culture, islet, acinar, and duct cells were used for βGAL analysis by FACS, GSIS, and staining. Washing with PBS and centrifugation, the supernatant was removed and 500-1000 μL of TrypLE (gibco 12604-021) was added to islet, acinar, and duct cells. All three types of cells were immediately heated in a 37°C water bath for 10 minutes. During this time, they were vortexed for 10 seconds every 3 minutes, and the reaction was stopped immediately after adding with serum-containing medium on ice. Dispersed cells were stained with βGAL using kit (ENZ-KIT 130010) and cultured for a total of 3 hours. The percentage of βGAL-positive cells was extracted using FACS Aria and analyzed using FlowJo. Our FACS gating strategy can be seen in the Supplemental Figure 4A.

**Static GSIS**

Ten islets of equal size were hand-picked and plated onto a 24-well plate. After incubating 2.8 mM low glucose for 1 hour, the islets were then cultured in 2.8mM low glucose and 20.2 mM high glucose for 1 hour each. The supernatants were analyzed by insulin ELISA (Mercodia Insulin 10-1113-10). The DNA content of the remaining islets was measured using QIAGEN DNAeasy and normalized.

**Islet Perifusion**

Islets received were allowed to rest overnight in CMRL media prior to perifusion. On the morning of the perifusion, a fresh KRBH buffer with BSA was prepared in order to create the glucose solutions required for the procedure. The islets were removed from the incubator and fifty islets were selected per chamber that was run on the Peri-Lite Instrument from BIOREP. The islets were placed into the perifusion chambers with the PERI-BEAD solution and the chambers filled with the KRBH buffer before being attached to the instrument. The islets were flushed with a low glucose solution (2.8mM glucose) prior to the experiment. The islets were then treated with a low glucose solution (2.8mM glucose), a high glucose solution (20.2mM glucose), and a high glucose solution with IBMX (1.2mM). Between each high glucose treatment, the islets were treated with low glucose before ending with an extended low glucose treatment. Each channel of the perifusion was run at 150ul/minute. The perfusate was collected in 96 well Nunc plates, covered with an adhesive film, and stored at -20*C until an ELISA was performed to measure the insulin content from the perifusion.

**RT-qPCR**

The total RNA isolated with RNeasy Micro Kit (QIAGEN, 74004) was reverse transcribed (MultiScribe Reverse Transcripase, Thermo Fisher Science, 4308228). We used SYBR green to detect, and specific primers for p16 and p21 were used. Samples were normalized to TBP, and the comparative CT (threshold cycle) method was used to calculate gene expression levels. The sequence of each gene. TBP 5’-TGTGCACAGGAGCCAAGAGT-3’ (forward), 5’-ATTTTCTTGCTGCCAGTCTGG-3’ (reverse). p16 5’-CTTCGGCTGACTGGCTG-3’ (forward), 5’-GCCTCCGACCGTAACTATTC-3’ (reverse). p21 5’-TGTCACTGTCTTGTACCCTTG-3’ (forward), 5’-GCGTTTGGAGTGGTAGAAATC-3’ (reverse).

**Staining**

After dispersing into single cells using TrypLE and 10 min incubation followed by overnight culture, dispersed pancreas tissues (islets, acinar, and duct) were fixed with 10% formalin. The cells were permeabilized using 0.3% Triton X and blocked using 1:50 Normal Donkey Serum and incubated overnight at 4°C with Ki67 (ab15580/abcam, 1:500) or HMGB1 (ab79823/abcam, 1:400), respectively. The cells were incubated with a 1:200 AlexaFluor™ 594 anti-Rabbit secondary antibody (111-585-003/Jackson ImmunoResearch Laboratories). Islets were additionally incubated with 1:25 Guinea Pig anti-Pig Insulin (PA1-29383/Invitrogen) and 1:200 AlexaFluor™ 488 anti-Guinea Pig secondary antibody (706-545-148/Jackson ImmunoResearch Laboratories). All cells were mounted with DAPI for nuclear staining (F6057/Sigma-Aldrich), and pictures were taken in the same setting with Zeiss confocal microscope.

**Bioinformatic analysis of scRNAseq**

The raw fastq files were inputted into Cell Ranger to obtain gene counts. CellBender was applied to eliminate ambient RNA or random barcode swapping counts, scDblFinder [6] to remove doublets, and a Seurat object was constructed. Using Seurat [7], only cells with more than 1000 RNA counts, more than 500 features, and less than 20% mitochondrial gene expression were kept, and normalized using the LogNormalize function, which calculates the natural logarithm of the counts plus one pseudocount. Each gene was scaled to have mean zero and variance of one. The principal components (PCs) were calculated and the top 114 were kept, since that was found to be the optimal number from JackStraw analysis. Using these PCs, the data were harmonized across donors using the R package Harmony [8]. K nearest neighbors were applied and clustered the cells using the Leiden algorithm, which yielded 19 clusters, and plotted as a UMAP. Clusters were defined based on marker genes and defined populations using *CDKN1A* and *CDKN2A*. Differential expression of cell populations utilized Seurat’s FindMarkers function and differential expression of samples at the pseudo-bulk level utilized Limma-Voom [9, 10]. Association was tested of variables such as senescent markers with gene expression at the pseudo-bulk level using Limma-Voom and donor effects were accounted for using the Limma function duplicateCorrelation, which is an analog of a mixed model [11]. Gene expression was compared across quartiles at the pseudo-bulk level accounting for donor effect with duplicate correlation and one-sided p-values were calculated. Senescence scores were assessed for each sample, cell type, and gene set using single-sample Gene Set Enrichment Analysis (ssGSEA), and the association was tested between the senescence scores and age or HbA1c using mixed effect linear regression with gender and BMI as fixed effect covariates, and subject as random slopes with the R package lmerTest [12]. Cell-cell interaction analysis used the R package CellChat [13].

**Bioinformatic analysis of Visium**

Spots exhibiting fewer than 1000 total transcript counts, fewer than 500 detected genes, or with more than 5% of total reads mapping to mitochondrial genes were excluded. Using Seurat, gene expression was normalized using the LogNormalize function, each gene was scaled to have mean zero and variance of one, the principal components (PCs) were calculated, and the top 77 were kept, since that was found to be the optimal number from JackStraw analysis. Using these PCs, the data were harmonized across donors using the R package Harmony [8]. K nearest neighbors was applied, the spots were clustered using the Leiden algorithm, and they were plotted as a UMAP. The cluster labels were projected back onto the original tissue images using Seurat’s SpatialDimPlot function. The cell type labels were transferred from the reference scRNAseq onto the Visium spots using Seurat and they were confirmed using the expression of marker genes. Spots were defined as having high or low levels of senescence markers using density estimates for each gene’s expression using a Gaussian finite mixture model from R package Mclust [14]. Differential expression or association of expression to age, BMI, HbA1c, or senescent markers were tested at the pseudo-bulk level with Limma-Voom while accounting for donor effects using Limma’s duplicateCorrelation. To test association between senotypes and immune cell infiltration in islets, islets were segmented from exocrine pancreas tissue using the R package Semla [15] and the percentages were calculated of *CDKN1A*-high, *CDKN2A*-high, and immune-high cells per islet. Arcsine square-root transformed was done to stabilize variance and improve normality and association tested using mixed models (with glmmTMB [16]) since there was nesting of cells within islets within samples within donors.

**Bioinformatic analysis of Xenium and CODEX**

Cells were removed with fewer than 50 total RNA counts, fewer than 30 detected genes, fewer than 30 total protein intensity units, or fewer than 10 detected proteins. The Xenium data were preprocessed by normalizing with LogNormalize, each feature scaled to have mean zero and variance of one, and PCA performed. The Codex data were normalized with Seurat’s centered log ratio, with scaling of each gene to have mean zero and variance of one, and PCA was applied. The optimal number of PCs were separately identified to be retained using the findPC package, which showed both should have 12 PCs. Harmony was run separately on both data modalities setting the batch variable as donor, as above. To integrate both data types, Seurat’s FindMultiModalNeighbors function was applied to compute a joint neighbor graph that integrates information from both the Xenium RNA PCs and the Codex protein PCs using a randomly sampled subset of the cells, as using all of the approximately 13 million cells would be computationally prohibitive. This graph was used to calculate UMAP embeddings. The non-sampled cells were projected onto this by calculating their nearest neighbors with the RcppHNSW package, which implements a hierarchical navigable small-world (HNSW) graph. As above, for RNA differential expression or association of expression to age, BMI, HbA1c, or senescent markers was tested at the pseudo-bulk level with Limma-Voom while accounting for donor effects using Limma’s duplicateCorrelation, whereas for proteins Limma was used without the Voom transformation. Quartile analysis was applied as with scRNAseq and Visium to p16, p21, and HMGB1 proteins, but not to CDKN1A and CDKN2A as these genes had too many zero values. Association was tested between senotypes and immune cell infiltration in islets as with the Visium data.

**Data and Code Availability**

Visium, scRNA-seq, Phenocycler, Xenium and H&E raw datasets are made available for the public via the SenNet Consortium portal (https://data.sennetconsortium.org/). Code associated with processing and analysis will be made available via the SenNet GitHub page (<https://github.com/sennetconsortium>).
