## Supplemental Figures for "Distinct senescent β-cell senotypes differentially drive islet aging and dysfunction"

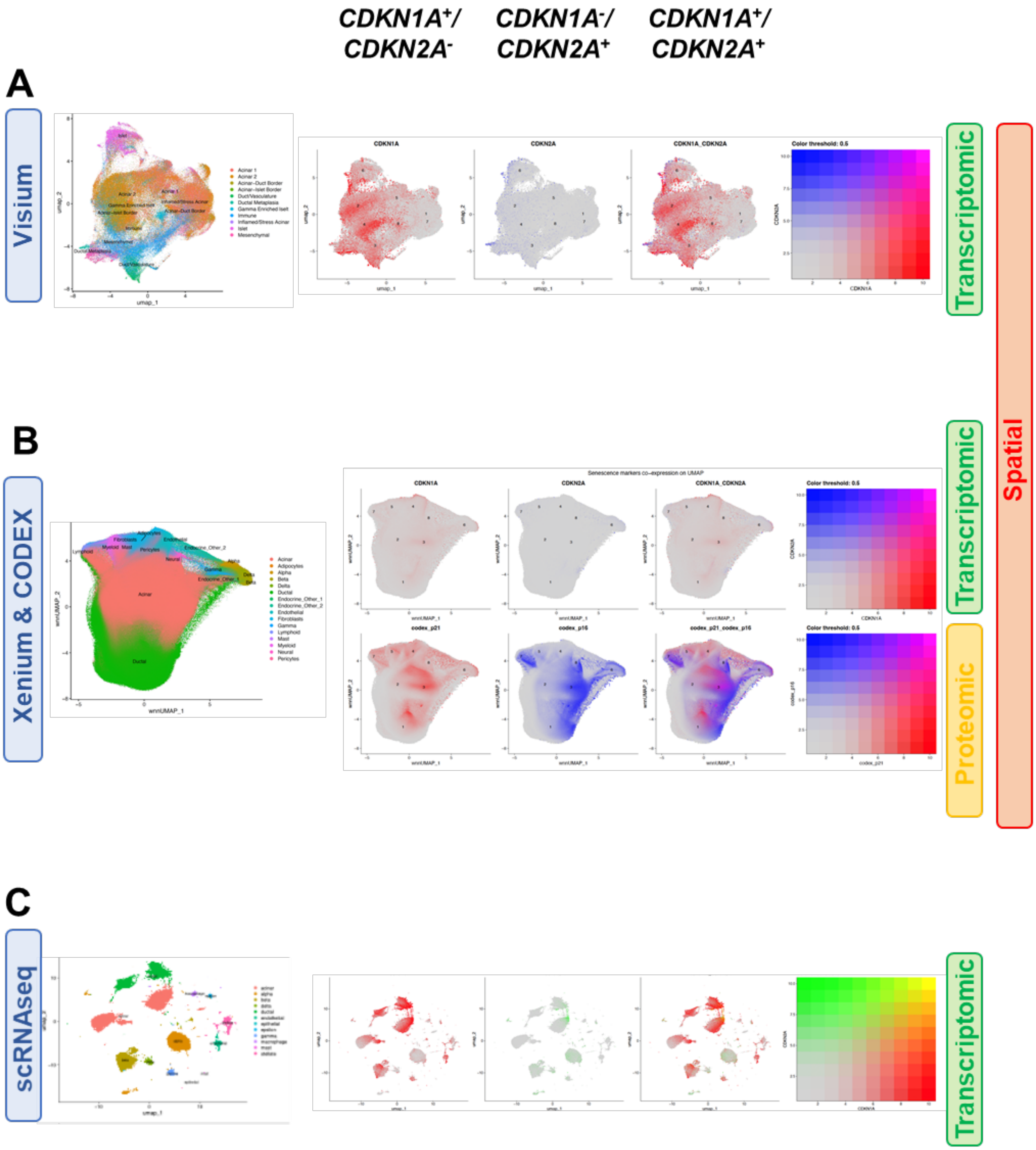


**Supplementary Figure 1 (Relates to Fig. 1).** UMAP of cell identity and senotypes in whole pancreas as detected by Visium (**A**) and integrated Xenium and CODEX (**B**). UMAP of dispersed pancreas (islet, acinar, and duct cells) and its senotypes analyzed in scRNAseq (**C**). See suppl. Table 1 for donor and sample numbers

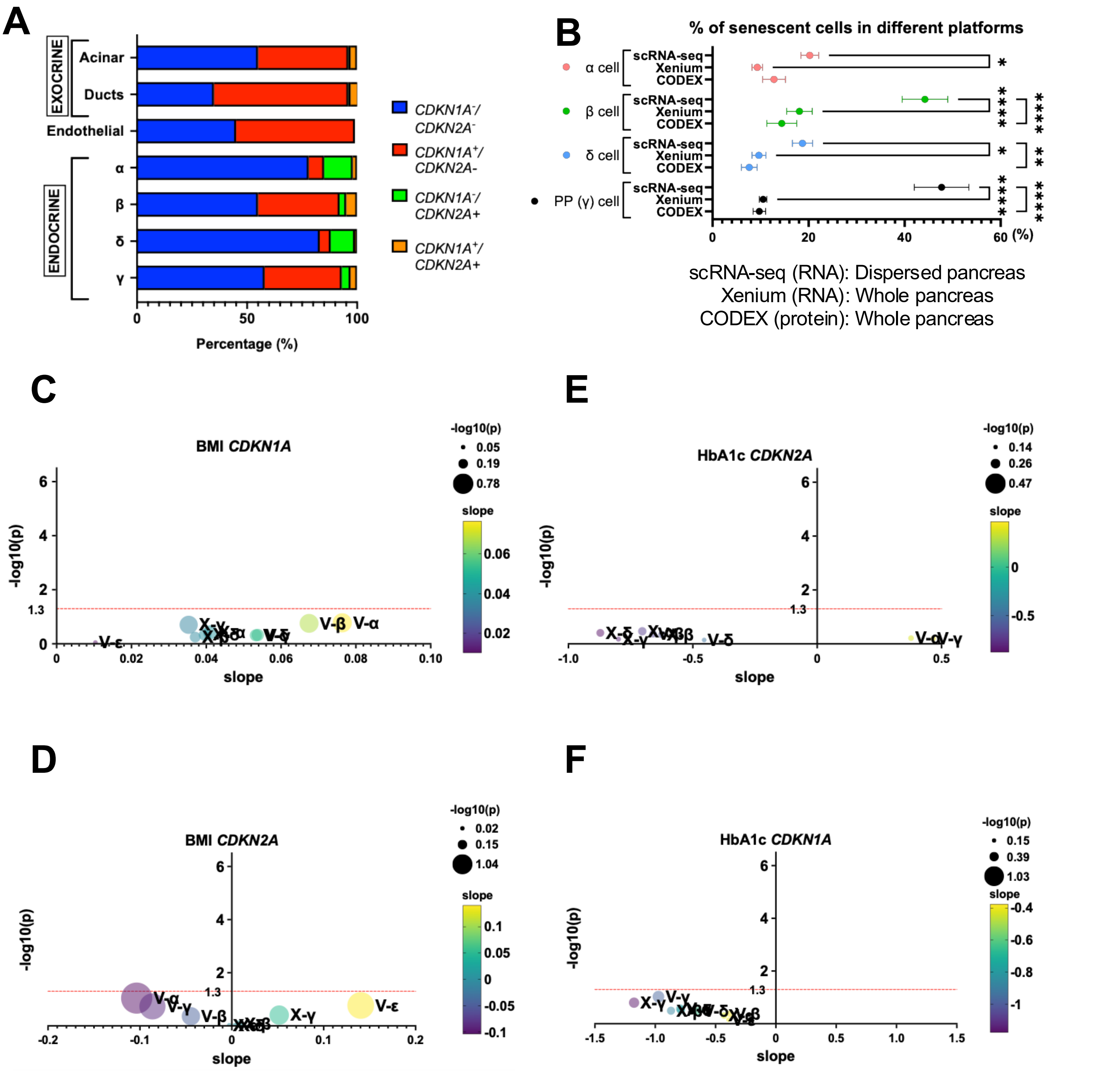


**Supplementary Figure 2 (Relates to Fig. 1).** (**A**) Percentage of senotypes (*CDKN1A*$^{+}$/*CDKN2A*$^{-}$, *CDKN1A*$^{-}$/*CDKN2A*$^{+}$, *CDKN1A*$^{-}$/*CDKN2A*$^{-}$, and *CDKN1A*$^{+}$/*CDKN2A*$^{+}$) in human endocrine cell types from scRNAseq data obtained from 10 donors. (**B**) Comparison of percentage of senescent cells per islet cell type across platforms. Correlation analysis between BMI and *CDKN1A* (**C**) and *CDKN2A* (**D**), HbA1c and *CDKN1A*

(**E**), and *CDKN2A* (**F**) in different endocrine pancreatic cell types (α, β, δ, γ and ε cells) from Visium (V) and Xenium (X) data.

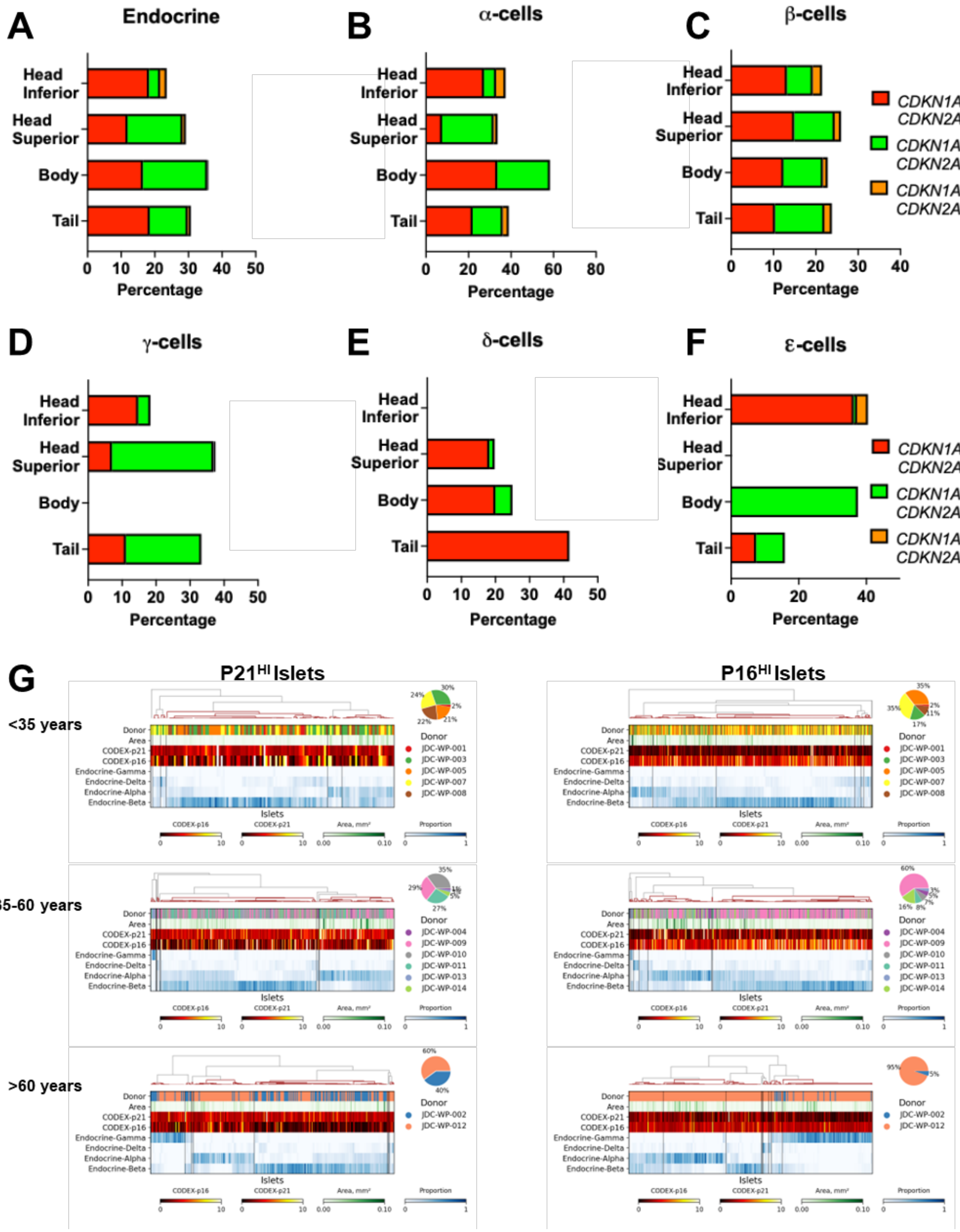

**Supplementary Figure 3. (Relates to Fig. 3)** (**A-F**) Senotype proportions of whole pancreas in 4 regions (Head Superior, Head Inferior, Body, and Tail) per pancreatic endocrine cell type. The average percentage of SnCs in α-cells in the pancreatic body varies widely among 10 donors (0-100%); (**G**) Hierarchical clustering of pancreatic islets reveals donor-specific endocrine composition patterns across different age groups and selection criteria. Islets from young (<35, n=5), middle-aged (35-60, n=6), and older donors (>60, n=2) were selected based on high expression of CODEX-p16 or CODEX-p21 (normalized fluorescence intensity >2.0) and hierarchically clustered using Pearson correlation with average linkage based on the proportions of α, β, δ, γ and ε cells. Each column represents an individual islet, with the top row indicating donor identity (color-coded) and the islet cross-sectional area shown in the green heatmap row. CODEX-p16 and CODEX-p21 expression levels are displayed as heatmaps (yellow = high, black = low, red = intermediate), while endocrine cell type proportions are shown in blue gradients (dark = high proportion). The dendrogram illustrates hierarchical relationships among islets, and pie charts show the proportion of islets contributed by each donor. The results demonstrate that islets cluster primarily by endocrine composition rather than by donor, with significant heterogeneity in senescence marker expression and cell type distribution across all age groups and selection objectives. The number of islets analyzed for each age range was as follows: <35 years- 520 islets; 35-60 years- 854 islets; >60 years- 561 islets.

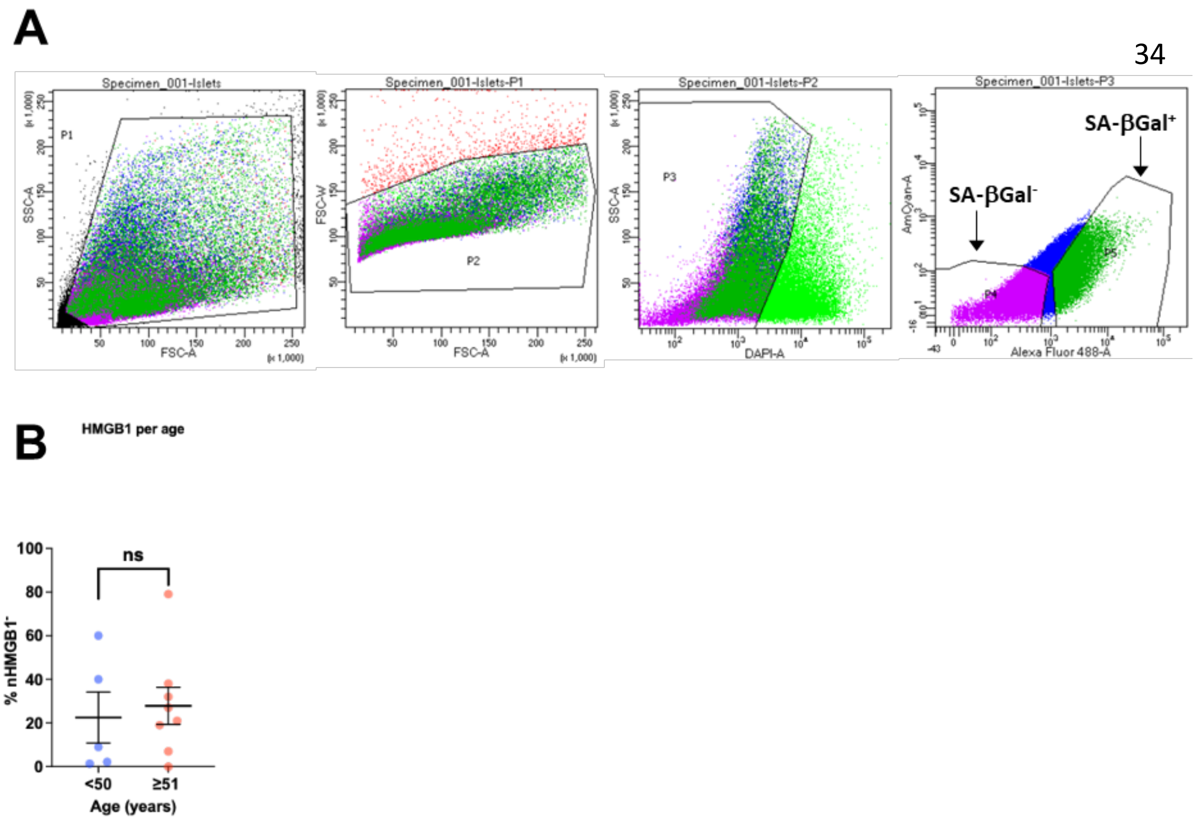


**Supplementary Figure 4 (Relates to Fig. 4).** (**A**) FACS sorting strategies for β-cells. Gating according to cell size (pulse width) to exclude doublets and triplets; Gating strategy based on granularity to exclude cell debris.; Gating strategy to select for DAPI- live cell population. After excluding dead cells with DAPI, the image was gated with SA-βGal. (**B**) Immunostaining quantification of HMGB1 staining in dispersed pancreases in donors of different ages.

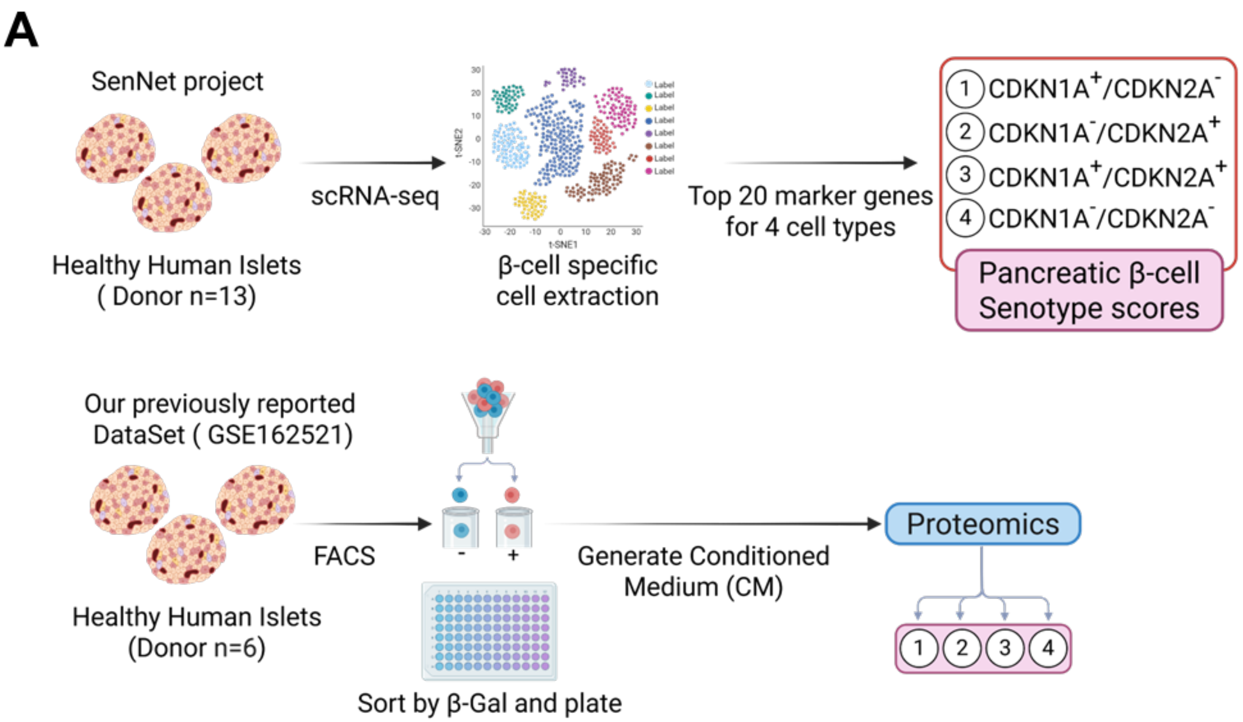


**Supplementary Figure 5 (Relates to Fig. 4).** (**A**) Graphical abstract of extracting top 20 marker genes in 3 cell types (*CDKN1A*$^{+}$*/CDKN2A*$^{-}$, *CDKN1A*$^{-}$*/CDKN2A*$^{+}$, *CDKN1A*$^{+}$*/CDKN2A*$^{+}$, *CDKN1A*$^{-}$*/CDKN2A*$^{-}$) in pancreatic β-cells (Top row). These markers were used using our previously reported datasets from proteomic data defining human islet SASP (GSE150285) [1]. of interactions or the strength of the communication.

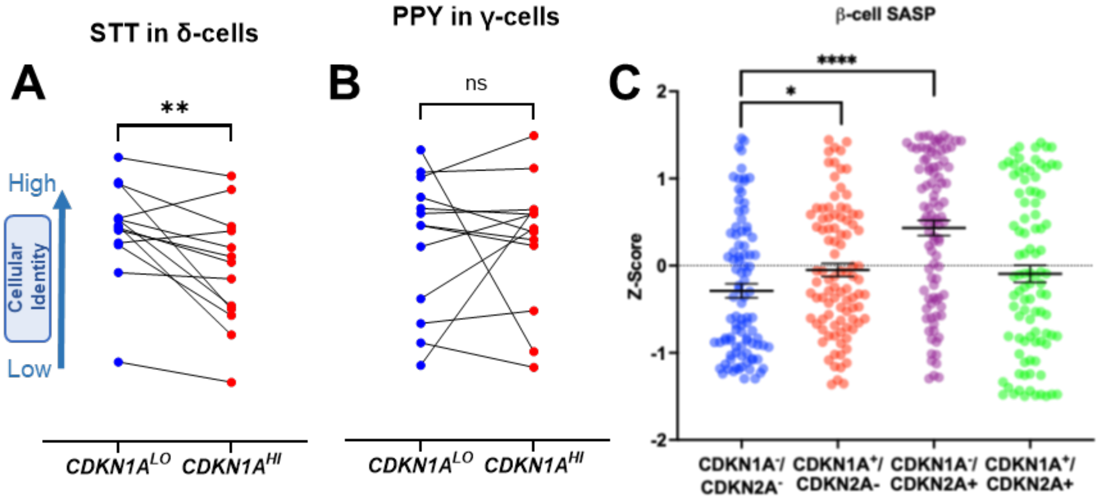


**Supplementary Figure 6 (Relates to Fig. 5).** (**A**) scRNASeq data per donor STT and (**B**) PPY expression across levels of *CDKN1A* in δ or γ cells. High and low expression of *CDKN1A*$^{HI}$ and *CDKN1A*$^{LO}$, n=13, Mean+/-SEM, paired t-test. (**C**) Gene expression Z-score scatter plot of SASP genes in β-cell senotypes.
