## Supplemental Tables for "Distinct senescent β-cell senotypes differentially drive islet aging and dysfunction"

**Supplemental Table 1. Donor Information**

| Whole Pancreas | | | | | | | Pancreas region | | |
|---|---|---|---|---|---|---|---|---|---|
| **Donor ID** | **Age** | **Sex** | **BMI** | **HbA1c (%)** | **Source** | **Specimen** | **Visium** | **Xenium + CODEX** | **iCLAP** |
| JDC_WP_01 | 21 | M | 28 | 4.9 | UTHSCSA | Whole Pancreas | HI/HS/B/T | HS/T | T |
| JDC_WP_02 | 71 | F | 26 | 5.4 | UTHSCSA | Whole Pancreas | HI/HS/B/T | HS/T | |
| JDC_WP_03 | 32 | M | 28 | 5.1 | UTHSCSA | Whole Pancreas | | HS/T | |
| JDC_WP_04 | 54 | M | 23.9 | 5.2 | UTHSCSA | Whole Pancreas | HI/HS/B/T | HS/T | |
| JDC_WP_05 | 28 | F | 17.7 | 5.3 | UTHSCSA | Whole Pancreas | HI/HS/B/T | HS/T | |
| JDC_WP_07 | 22 | M | 22.4 | 5.1 | UTHSCSA | Whole Pancreas | HI/HS/B/T | HS/T | |
| JDC_WP_08 | 23 | F | 17.2 | N/A | UTHSCSA | Whole Pancreas | HI/HS/B/T | HI/T | |
| JDC_WP_09 | 40 | F | 23.0 | 5.2 | UTHSCSA | Whole Pancreas | HI/HS/B/T | HS/T | |
| JDC_WP_10 | 37 | F | 23.5 | 5.9 | UTHSCSA | Whole Pancreas | HI/HS/B/T | HS/T | |
| JDC_WP_11 | 47 | M | 21.8 | 4.8 | UTHSCSA | Whole Pancreas | HI/HS/B/T | HS/T | |
| JDC_WP_12 | 69 | F | 18 | N/A | UTHSCSA | Whole Pancreas | HI/HS/B/T | HI/HS/T | |
| JDC_WP_13 | 35 | M | N/A | 5.4 | UTHSCSA | Whole Pancreas | | HS/T | |
| JDC_WP_14 | 39 | M | 24.7 | N/A | UTHSCSA | Whole Pancreas | | HS/T | |
| JDC_WP_15 | 43 | M | 14.4 | 5.8 | UTHSCSA | Whole Pancreas | | HS/T *Xenium | |
| JDC_WP_16 | 43 | M | 23.3 | N/A | UTHSCSA | Whole Pancreas | | HS/T *Xenium | |
| JDC_WP_26 | 82 | F | 26 | 5.5 | Imagine Pharma | Whole Pancreas | | | T |

**HI: Head Inferior, HS: Head Superior, B: Body, T: Tail**

| Dispersed Pancreas | | | | | | | |
|---|---|---|---|---|---|---|---|
| **Donor ID** | **Age** | **Sex** | **BMI** | **HbA1c (%)** | **Source** | **Specimen** | **scRNA-seq** |
| PLBD001PAD | 46 | M | 23.1 | 4.9 | Prodo Labs | Dispersed Pancreas | islet/acinar/duct |
| PLBD002PAD | 42 | F | 29.3 | 5.2 | Prodo Labs | Dispersed Pancreas | islet/acinar/duct |
| PLBD003PAD | 50 | F | 28.3 | 5.4 | Prodo Labs | Dispersed Pancreas | islet/acinar/duct |
| PLBD004PAD | 57 | M | 30.9 | 5.2 | Prodo Labs | Dispersed Pancreas | islet/acinar/duct |
| PLBD005PAD | 59 | M | 24.5 | 5.2 | Prodo Labs | Dispersed Pancreas | islet/acinar/duct |
| PLBD006PAD | 58 | F | 31 | 4.7 | Prodo Labs | Dispersed Pancreas | islet/acinar/duct |
| PLBD007PAD | 64 | M | 25.5 | 5.2 | Prodo Labs | Dispersed Pancreas | islet/acinar/duct |
| PLBD008PAD | 47 | M | 28.3 | 5.3 | Prodo Labs | Dispersed Pancreas | islet/acinar/duct |
| PLBD009PAD | 57 | F | 30.2 | 5.8 | Prodo Labs | Dispersed Pancreas | islet/acinar/duct |
| PLBD010PAD | 58 | M | 28.9 | 5.5 | Prodo Labs | Dispersed Pancreas | islet/acinar/duct |
| PLBD011PAD | 34 | M | 24.6 | 5.4 | Prodo Labs | Dispersed Pancreas | islet/acinar/duct |
| PLBD012PAD | 61 | M | 29.5 | 5.7 | Prodo Labs | Dispersed Pancreas | islet/acinar/duct |
| PLBD013PAD | 69 | M | 29.4 | 5.4 | Prodo Labs | Dispersed Pancreas | islet/acinar/duct |

| | **Donor** | **Sample** |
|---|---|---|
| Xenium | 10 | 32 |
| Visium | 10 | 40 |
| CODEX | 10 | 32 |
| scRNAseq | 13 | 42 |
| Proteomics | 9 | 17 |
| iCLAP protein | 2 | |
| Function | 11 | |

**Supplemental Table 2. Senescence Signatures**

| α (Alpha) | β (Beta) | δ (Delta) | γ (Gamma) |
|---|---|---|---|
| HRAS | PML | SIRT1 | HBS1L |
| PRKDC | SRF | NRG1 | CCN2 |
| HPS5 | ZMPSTE24 | KAT6A | RHOB |
| CDKN2B | PLA2R1 | CLTB | MAP1LC3B |
| HSPA2 | IRF7 | IRF7 | RAB31 |
| NRG1 | SERPINE1 | CDKN2B | EIF2S2 |
| PEA15 | S100A11 | PAWR | PEA15 |
| RAB31 | TBX2 | PNPT1 | MIF |
| PAWR | NUAK1 | IGFBP3 | RAB13 |
| MMP1 | NRG1 | MTOR | GUK1 |
| SMURF2 | NSMCE2 | RAB5B | PNPT1 |
| S100A11 | PLK2 | MAP3K3 | B2M |
| TES | NEK4 | HBS1L | BCL6 |
| SERPINE1 | NUP62 | RAB31 | CALR |
| OPA1 | RAB5B | CDKN2A | IGFBP4 |
| PML | CDKN2B | FBXO5 | VASH1 |
| SIRT6 | TP53 | MDM2 | RRAS |
| NSMCE2 | RABGGTA | ING1 | CREG1 |
| MDM2 | YPEL3 | CDKN1A | IFI16 |
| FN1 | MTOR | CDK6 | TES |
| SPARC | CDKN2A | SRF | IRF5 |
| CCN2 | IRF5 | EIF2S2 | KRAS |
| STAT1 | PRELP | RBL1 | OPTN |
| CDKN1A | MAP1LC3B | PML | CDKN2B |
| TNFAIP3 | STAT1 | DNAJA3 | RAC1 |
| THBS1 | BCL2L2 | TOP2B | CDKN2A |
| IGFBP1 | BRCA2 | PLK2 | CDKN1A |
| PRKCD | IGFBP7 | HRAS | ZMPSTE24 |
| TP53 | CCN2 | TP53 | NUP62 |
| TERF2 | CDKN1A | S100A11 | NEK4 |
| LMNA | PRMT6 | BMPR1A | HTT |
| IRF7 | TSPYL5 | TERF2 | FILIP1L |
| TNFAIP2 | ING1 | ZMPSTE24 | OPA1 |
| VIM | FN1 | SUV39H1 | RGL2 |
| IGSF3 | ID2 | SIRT6 | ICMT |
| ECRG4 | CCND1 | ZKSCAN3 | IRF7 |
| CDKN2A | MDM2 | PEA15 | TNFAIP3 |
| CLTB | RBL2 | MAP1LC3B | HSPA2 |
| CDKN2D | TES | VIM | F3 |
| OPTN | ALDH1A3 | MAPKAPK5 | NME2 |
| FILIP1L | IGFBP5 | CDKN2D | S100A11 |

| | | | |
|---|---|---|---|
| CCND1 | F3 | HSPA2 | MDM2 |
| PLA2R1 | CDKN2D | GSN | CDK6 |
| MAP3K3 | KRAS | SMC6 | SUV39H1 |
| SPI1 | CALR | NSMCE2 | CLTB |
| FBXO5 | CLTB | NEK4 | MAPKAPK5 |
| KRAS | CREG1 | PRMT6 | HMGA1 |
| RABGGTA | FBXO5 | TES | PAWR |
| BRCA2 | HTATIP2 | YPEL3 | LMNA |
| RSL1D1 | TGFB1I1 | RABGGTA | SMPD1 |
| MTOR | RSL1D1 | RSL1D1 | IGFBP6 |
| HTATIP2 | RHOB | NAMPT | MAP2K3 |
| CGAS | MAP2K1 | CD44 | ID2 |
| CDK6 | CGAS | HPS5 | PTEN |
| EEF1E1 | DNAJA3 | PRKCD | TOP2B |
| FZR1 | GSN | ID2 | HTATIP2 |
| HMGA1 | SPI1 | LMNA | FBXO4 |
| ING1 | HTT | HTT | MTOR |
| PNPT1 | MMP1 | ICMT | TERF2 |
| SIRT1 | HMGA1 | FZR1 | PRKCD |
| ZMIZ1 | GCH1 | CDKN1C | ZMIZ1 |
| TGFB1I1 | IFI16 | PTEN | TP53 |
| DNAJA3 | SMURF2 | HTATIP2 | ISG15 |
| ID2 | HRAS | OPA1 | BMPR1A |
| CRYAB | SMPD1 | NDN | STAT1 |
| MAP2K3 | CD44 | TNFAIP2 | KAT6A |
| IRF5 | VIM | MAP2K3 | ING1 |
| ISG15 | SUV39H1 | IGFBP5 | ARG2 |
| BCL6 | GUK1 | MORC3 | PML |
| SUV39H1 | PRKCD | ARG2 | PLK2 |
| GSN | THBS1 | EEF1E1 | EEF1E1 |
| RHOB | ISG15 | F3 | RBL2 |
| PLK2 | BCL6 | STAT1 | NSMCE2 |
| EIF2S2 | FILIP1L | NUAK1 | NUAK1 |
| RBL1 | CRYAB | RGL2 | FZR1 |
| SRF | ZKSCAN3 | OPTN | RBL1 |
| YBX1 | LMNA | BCL2L2 | IGFBP2 |
| RAB5B | FBXO4 | SIRT1 | SOD1 |
| IGFBP4 | IGFBP6 | | CD44 |
| RAC1 | HBS1L | | CDKN2D |
| ZMPSTE24 | PAWR | | ILK |
| ZKSCAN3 | PRKDC | | BCL2L2 |
| MORC3 | RAC1 | | NAMPT |
| ILK | TNFAIP2 | | MAP3K3 |

| | | | |
|---|---|---|---|
| RRAS | CDKN1C | | FBXO5 |
| MAP1LC3B | EEF1E1 | | WRN |
| ABL1 | RAB31 | | YPEL3 |
| NAMPT | ICMT | | TSPYL5 |
| NDN | MAP3K3 | | RSL1D1 |
| NME2 | SIRT1 | | HRAS |
| BMPR1A | NME2 | | ZKSCAN3 |
| PRMT6 | ILK | | YBX1 |
| YPEL3 | IGSF3 | | IGFBP5 |
| ARG2 | IGFBP4 | | SMURF2 |
| WRN | RBL1 | | PRKDC |
| SMC5 | PEA15 | | |
| | OPA1 | | |
| | TNFAIP3 | | |
| | COL1A2 | | |
| | ECRG4 | | |

**Supplemental Table 3. SASP Signatures**

| α (Alpha) | β (Beta) | δ (Delta) | γ (Gamma) |
|---|---|---|---|
| ITGA2 | SERPINE1 | BMP2 | TNFRSF10C |
| NRG1 | TGFB1 | NRG1 | MIF |
| SEMA3F | SPP1 | ANG | IGFBP4 |
| PLAU | ETS2 | IQGAP2 | C3 |
| SELPLG | NRG1 | VEGFC | ETS2 |
| MMP9 | FGF2 | ITGA2 | FGF1 |
| CSF1 | SCAMP4 | TNFRSF10C | TGFB1 |
| TGFB1 | MMP2 | IGFBP3 | CXCL2 |
| TNFRSF11B | TNFRSF11B | CXCL16 | GDF15 |
| SERPINE1 | TNFRSF10C | CSF1 | SERPINE2 |
| EGFR | IQGAP2 | GEM | CXCL3 |
| FAS | FGF7 | FAS | BMP6 |
| BMP2 | FAS | PLAUR | ANG |
| CD9 | IL18 | IL7 | CTNNB1 |
| IL7 | IL32 | SCAMP4 | CX3CL1 |
| IL13 | INHBA | GDF15 | BMP2 |
| GDF15 | CCL18 | TGFB1 | ACVR1B |
| CCL2 | GDF15 | CTNNB1 | ICAM1 |
| TNFRSF10C | CTSB | ICAM1 | VGF |
| IGFBP1 | PECAM1 | TNFRSF11B | CST3 |
| MMP14 | IGFBP7 | INHBA | ITGA2 |
| TNFRSF1A | GEM | C3 | PLAUR |
| CXCL12 | AREG | IL6ST | IGFBP6 |
| CRP | CTNNB1 | TN4SF1 | IL7 |
| AREG | EGFR | VGF | CXCL8 |
| PGF | HGF | NAP1L4 | CTSB |
| EDN1 | CX3CL1 | ACVR1B | CSF1 |
| CXCL3 | PLAU | JUN | BEX3 |
| IL15 | CXCL16 | PTBP1 | IL15 |
| TN4SF1 | CRP | IGFBP5 | CXCL16 |
| INHBA | CXCL3 | TIMP2 | INHA |
| PLAUR | CSF1 | BEX3 | JUN |
| C3 | CD9 | CTSB | SPP1 |
| ICAM3 | LCP1 | | GEM |
| BMP6 | IGFBP5 | | CD9 |
| IL32 | ANG | | TNFSF10 |
| VEGFC | PAPPA | | TUBGCP2 |
| ETS2 | CD55 | | IGFBP2 |
| SPP1 | MPO | | VEGFC |
| CD55 | IL15 | | VEGFA |
| ICAM1 | CXCL12 | | SCAMP4 |

| | | | |
|---|---|---|---|
| IQGAP2 | CXCL1 | | INHBA |
| CTNNB1 | PGF | | PIGF |
| CXCL2 | PTBP1 | | IL6ST |
| JUN | CCL2 | | TM4SF1 |
| GEM | SELPLG | | |
| ACVR1B | GPNMB | | |
| CXCL8 | SERPINE2 | | |
| IGFBP4 | EDN1 | | |
| ANGPTL4 | C3 | | |
| CX3CL1 | ANGPTL4 | | |
| PTBP1 | TM4SF1 | | |
| IL6ST | TNFRSF1A | | |
| CXCL1 | MMP14 | | |
| NAP1L4 | TIMP2 | | |
| CXCL16 | SOST | | |
| TIMP2 | CXCL2 | | |
| VEGFA | VGF | | |
| IGFBP6 | VEGFC | | |
| | IL7 | | |
| | ITGA2 | | |
| | IGFBP6 | | |
| | SEMA3F | | |
| | INHA | | |
| | BMP2 | | |
| | IL6ST | | |
| | CXCL8 | | |
| | PLAUR | | |
| | IGFBP4 | | |
